# A neuron-specific microexon ablates the novel DNA-binding function of a histone H3K4me0 reader PHF21A

**DOI:** 10.1101/2023.10.20.563357

**Authors:** Robert S. Porter, Masayoshi Nagai, Sojin An, Maria C. Gavilan, Yumie Murata-Nakamura, Katherine M. Bonefas, Bo Zhou, Olivier Dionne, Jeru Manoj Manuel, Joannie St-Germain, Liam Browning, Benoit Laurent, Uhn-Soo Cho, Shigeki Iwase

**Affiliations:** Department of Human Genetics, University of Michigan Medical School, Ann Arbor, MI 48109, USA; Department of Biological Chemistry, University of Michigan Medical School, Ann Arbor, MI 48109, USA; Genetics and Genomics Graduate Program, University of Michigan Medical School, Ann Arbor, MI 48109, USA. Michigan Neuroscience Institute, University of Michigan, Ann Arbor, MI 48109, USA; Neuroscience Graduate Program, University of Michigan Medical School, Ann Arbor, MI 48109, USA; Research Center on Aging, Centre Intégré Universitaire de Santé et Services Sociaux de l’Estrie-Centre Hospitalier Universitaire de Sherbrooke, Sherbrooke, QC, Canada; Department of Biochemistry and Functional Genomics, Faculty of Medicine and Health Sciences, Université de Sherbrooke, Sherbrooke, QC, Canada; Department of Pediatrics, University of Michigan Medical School, Ann Arbor, MI 48109, USA; Michigan Neuroscience Institute, University of Michigan, Ann Arbor, MI 48109, USA

**Author notes:** Department of Neurology, Mass General Brigham, Boston, MA, 02114, USA.

## Abstract

How cell-type-specific chromatin landscapes emerge and progress during metazoan ontogenesis remains an important question. Transcription factors are expressed in a cell-type-specific manner and recruit chromatin-regulatory machinery to specific genomic loci. In contrast, chromatin-regulatory proteins are expressed broadly and are assumed to exert the same intrinsic function across cell types. However, human genetics studies have revealed an unexpected vulnerability of neurodevelopment to chromatin factor mutations with unknown mechanisms. Here, we report that 14 chromatin regulators undergo evolutionary-conserved neuron-specific splicing events involving microexons. Of the 14 chromatin regulators, two are integral components of a histone H3K4 demethylase complex; the catalytic subunit LSD1 and an H3K4me0-reader protein PHF21A adopt neuron-specific forms. We found that canonical PHF21A (PHF21A-c) binds to DNA by AT-hook motif, and the neuronal counterpart PHF21A-n lacks this DNA-binding function yet maintains H3K4me0 recognition intact. In-vitro reconstitution of the canonical and neuronal PHF21A-LSD1 complexes identified the neuronal complex as a hypomorphic H3K4 demethylating machinery with reduced nucleosome engagement. Furthermore, an autism-associated *PHF21A* missense mutation, 1285 G>A, at the last nucleotide of the common exon immediately upstream of the neuronal microexon led to impaired splicing of *PHF21A*-n. Thus, ubiquitous chromatin regulatory complexes exert unique intrinsic functions in neurons via alternative splicing of their subunits and potentially contribute to faithful human brain development.

## Introduction

A long-standing question in biology is how cell-type-specific transcription programs emerge and progress during metazoan development. DNA-binding transcription factors (TFs) play crucial roles in this process. TFs are often expressed in specific cell types, bind to their cognate DNA sequences at promoters and enhancers, and direct transcription at specific genomic loci^1,2^. In multicellular organisms, DNA is organized into nucleosomes with the four core histones, making DNA refractory to the actions of RNA polymerase II^3^. To relax the inherently closed structure, chromatin regulators, including histone-modifying enzymes and chromatin-remodeling complexes, are recruited with a variety of mechanisms, including cell-type-specific TFs^4,5^. Unlike TFs, most chromatin regulators are expressed ubiquitously. A widely held assumption is that chromatin regulators exert the same intrinsic activities across cell types, and TFs provide the locus specificity.

The exceptions to the ubiquitous presence of chromatin regulatory machinery have been described, in which cell-type-specific subunits, encoded by separate genes, can direct unique transcription programs. Brg/Brm-associated factor (BAF) complex, an ATP-dependent chromatin-remodeling machine, carries BAF53b, BAF45b, and BAF45c in the neuronal lineage^6,7^, while an alternative complex carries BAF60c and BAF170 in the cardiac cell lineage^8,9^. Furthermore, the transcriptional preinitiation complex TFIID contains germ-cell-specific components such as TRF2, TRF3, and TAF7L during gametogenesis^10^. In addition, TFIID uniquely employs TAF9B subunit in neurons to promote transcription^11^. These complexes bearing cell-type-specific components play important roles in gene expression relevant to given tissue biology; however, the molecular functions of unique components within the complexes are not well understood.

Alternative mRNA splicing has emerged as another mechanism for modulating chromatin regulator functions. The brain transcriptomes exhibit more complex alternative splicing than other tissues^12–14^. The functional studies of splicing events have mainly concerned proteins relevant to cell biology unique to neurons such as synapses. However, several broadly-expressed chromatin regulators are known to undergo neuron-specific splicing events, such as TATA-binding protein-associated factor TAF1^15^, H3K9 methyltransferase G9a^16^, and histone H3 lysine demethylase LSD1^17^.

Among these regulators, LSD1, the first-identified H3K4me1/2 demethylase^18,19^, undergoes a well-characterized neuronal splicing event. An evolutionary-conserved 12-nucleotide exon encoding the four amino acids, Asp-Thr-Val-Lys, is inserted within the LSD1 catalytic domain uniquely in neurons^17^. Neuron-specific LSD1 isoform (herein referred to as LSD1-n) coexists with its canonical counterpart (LSD1-c), and the ratio between the two isoforms changes during neuronal differentiation as well as neuronal activation^20–22^. Notably, Thr369 within the neuronal exon can be phosphorylated. This phosphorylation changes local LSD1 conformation and reduces interaction with known corepressors, including HDAC1/2 and CoREST^23^.

However, the impact of LSD1 neuronal exon on its enzymatic activity remains controversial. The first report demonstrated that neuronal exon reduced its enzymatic activity on H3K4me without affecting the overall LSD1 structure^17^. The hypomorphic LSD1-n has been proposed to act as a dominant-negative form, which counteracts the H3K4me removal and transcriptional repression mediated by the LSD1-c^21,22^. Subsequent work found that the neuronal exon renders LSD1 new substrates—either H4K20me^24^ or H3K9me through cooperation with SVIL protein^20^. Despite the debatable impact on substrate specificity, ample genetic evidence indicates the importance of the LSD1 microexon in transcription, neuronal maturation, and cognitive behaviors^20–24^.

Aside from the TAF1, G9a, and LSD1, we do not know how prevalent the cell-type-specific alternative-splicing in chromatin regulators is. In this study, we performed an in-silico survey and identified 14 chromatin regulators that undergo evolutionary-conserved neuron-specific alternative splicing. We then determined the impact of two neuronal splicing events within the LSD1 complex, namely in PHF21A and LSD1. Furthermore, a human mutation associated with autism interfered with the production of the PHF21A-n form. Thus, alternative mRNA splicing can bestow unique chromatin-regulatory activity upon neurons for faithful brain development.

## Results

### Neuron-Specific Microexon Splicing of Chromatin Regulators

To systematically identify neuron-specific chromatin regulator isoforms, we intersected 801 human chromatin-regulatory proteins, defined in the Epifactors database^25^, with the list of neuron-specific alternative splicing events^12^. This analysis yielded 76 neuron-specific splicing candidates in chromatin regulators^26^. We then verified the neuron specificity of each candidate by manually inspecting the RNA-seq coverage of mouse brain cell types^27^ and human tissues (GTEx)^28^ with the UCSC genome browser. This filtering resulted in 16 conserved high-confidence neuron-specific alternative splicing events in 14 genes (Table 1), including LSD1 and TAF1. The G9a neuronal splicing^16^ was confirmed in the mouse datasets but not in humans. Most splicing events (13/14) lead to in-frame alternative polypeptides, which are relatively small (4 - 75 aa) and present in both conserved domains and unannotated protein segments. 7 splicing events (44%) represent microexons, encoding < 10 aa differences^12^. Only one event in KDM5B generates a stop codon, resulting in a truncated protein. In addition, two events occur as alternative transcription start sites, while the rest represent internal exons. The 14 molecules participate in multi-subunit machinery, such as BAF, MLL/COMPASS, PRC1, NuRD, and LSD1 complexes, all of which play critical roles in animal development. This survey also revealed two splicing events of ASH2L and HDAC7 that are not annotated in the GENCODE or Refseq transcript databases (Figure S1). Thus, a selected few broadly-expressed chromatin regulators can be spliced uniquely in neurons across species.

**Table 1:**
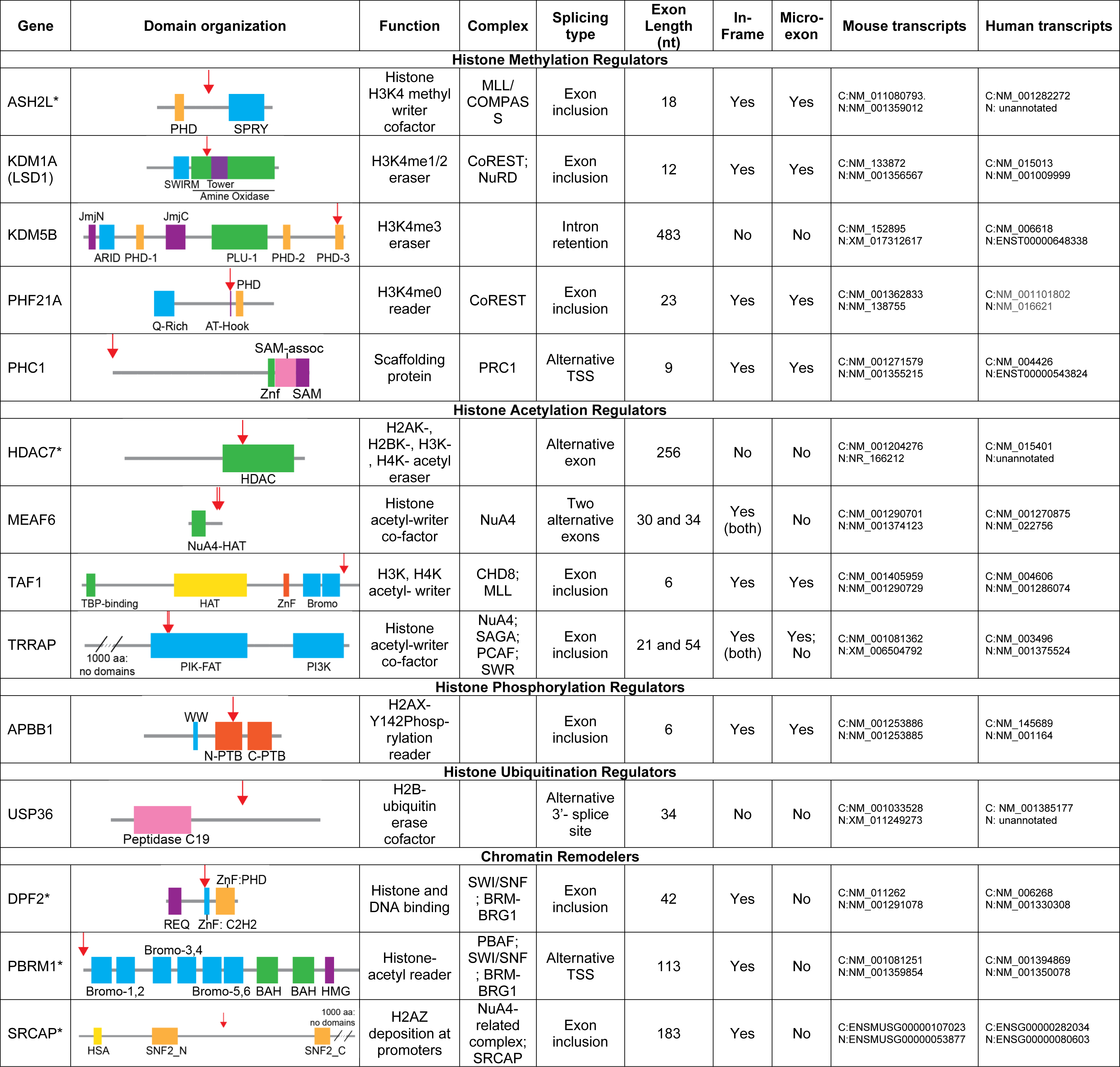
Evolutionary-conserved neuron-specific splicing events of chromatin regulators. WW: Two tryptophan domain that binds proline-rich peptide motifs. PTB: Phosphotyrosine-binding domain (N-terminal and C-terminal). REQ: Requiem-N-terminal domain of DPF2. ZnF: Zinc-Finger. C2H2: classic-type (Cys2-His2) zinc finger. PHD: plant homeodomain. HDAC: Histone Deacetylase domain. SWIRM: SWI3, RSC8, and MOIRA domain. Jmj: Jumonji histone demethylation domain (N-terminal and C-terminal). PLU-1: PLU-1 like protein. ARID: AT-rich DNA-interacting Domain. Q-Rich: Glutamine-rich region. AT-Hook: DNA-binding motif with a preference for A/T-rich regions. HAT: Histone acetyl-transferase. Bromo: Bromodomain. PIK-FAT: Phosphatidylinositol kinase-FRAP, ATM, and TRRAP subfamily. PI3K: Phosphatidylinositol 3-kinase. SPRY: Domain in the SPIa and the RYanodine receptor. NuA4 HAT: Histone acetyltransferase subunit NuA4. BAH: Bromo Adjacent Homology domain. HMG: High Mobility Group-box domain. HSA: Helicase/SANT-associated domain. SNF2: Sucrose Non-Fermenting family helicases (N and C terminal). SAM: Sterile Alpha Motif domain of Scm. Peptidase C19: peptidase family containing ubiquitinyl hydrolases. Asterisks (*) represent genes with splicing events observed in post-synaptic tissues, including skeletal and cardiac muscle tissues in addition to the brain.

| Gene | Domain organization | Function | Complex | Splicing type | Exon Length (nt) | In-Frame | Micro-exon | Mouse transcripts | Human transcripts |
| --- | --- | --- | --- | --- | --- | --- | --- | --- | --- |
| Histone Methylation Regulators |  |  |  |  |  |  |  |  |  |
| ASH2L* |  | Histone H3K4 methyl writer cofactor | MLL/COMPASS | Exon inclusion | 18 | Yes | Yes | C:NM_011080793.<br>N:NM_001359012 | C:NM_001282272<br>N: unannotated |
| KDM1A (LSD1) |  | H3K4me1/2 eraser | CoREST; NuRD | Exon inclusion | 12 | Yes | Yes | C:NM_133872<br>N:NM_001356567 | C:NM_015013<br>N:NM_001009999 |
| KDM5B |  | H3K4me3 eraser |  | Intron retention | 483 | No | No | C:NM_152895<br>N:XM_017312617 | C:NM_006618<br>N:ENST00000648338 |
| PHF21A |  | H3K4me0 reader | CoREST | Exon inclusion | 23 | Yes | Yes | C:NM_001362833<br>N:NM_138755 | C:NM_001101802<br>N:NM_016621 |
| PHC1 |  | Scaffolding protein | PRC1 | Alternative TSS | 9 | Yes | Yes | C:NM_001271579<br>N:NM_001355215 | C:NM_004426<br>N:ENST00000543824 |
| Histone Acetylation Regulators |  |  |  |  |  |  |  |  |  |
| HDAC7* |  | H2AK-, H2BK-, H3K-, H4K- acetyl eraser |  | Alternative exon | 256 | No | No | C:NM_001204276<br>N:NR_166212 | C:NM_015401<br>N:unannotated |
| MEAF6 |  | Histone acetyl-writer co-factor | NuA4 | Two alternative exons | 30 and 34 | Yes (both) | No | C:NM_001290701<br>N:NM_001374123 | C:NM_001270875<br>N:NM_022756 |
| TAF1 |  | H3K, H4K acetyl- writer | CHD8; MLL | Exon inclusion | 6 | Yes | Yes | C:NM_001405959<br>N:NM_001290729 | C:NM_004606<br>N:NM_001286074 |
| TRRAP |  | Histone acetyl-writer co-factor | NuA4; SAGA; PCAF; SWR | Exon inclusion | 21 and 54 | Yes (both) | Yes; No | C:NM_001081362<br>N:XM_006504792 | C:NM_003496<br>N:NM_001375524 |
| Histone Phosphorylation Regulators |  |  |  |  |  |  |  |  |  |
| APBB1 |  | H2AX-Y142Phosph-rylation reader |  | Exon inclusion | 6 | Yes | Yes | C:NM_001253886<br>N:NM_001253885 | C:NM_145689<br>N:NM_001164 |
| Histone Ubiquitination Regulators |  |  |  |  |  |  |  |  |  |
| USP36 |  | H2B-ubiquitin erase cofactor |  | Alternative 3'- splice site | 34 | No | No | C:NM_001033528<br>N:XM_011249273 | C: NM_001385177<br>N: unannotated |
| Chromatin Remodelers |  |  |  |  |  |  |  |  |  |
| DPF2* |  | Histone and DNA binding | SWI/SNF; BRM-BRG1 | Exon inclusion | 42 | Yes | No | C:NM_011262<br>N:NM_001291078 | C:NM_006268<br>N:NM_001330308 |
| PBRM1* |  | Histone-acetyl reader | PBAF; SWI/SNF; BRM-BRG1 | Alternative TSS | 113 | Yes | No | C:NM_001081251<br>N:NM_001359854 | C:NM_001394869<br>N:NM_001350078 |
| SRCAP* |  | H2AZ deposition at promoters | NuA4-related complex; SRCAP | Exon inclusion | 183 | Yes | No | C:ENSMUSG000000107023<br>N:ENSMUSG000000053877 | C:ENSG000000282034<br>N:ENSG000000080603 |

### Identification of PHF21A-n

Among the 14 factors, we focused on PHF21A (aka BHC80), an integral component of the canonical LSD1 complex^29,30^. Within the complex, PHF21A binds to unmethylated H3K4 (H3K4me0), the reaction product of LSD1, and further promotes the recruitment of LSD1 on chromatin^31^. In addition to PHF21A, the LSD1 complex harbors other stoichiometric components, including class I histone deacetylases HDAC1 and HDAC2, HMG-box DNA binding protein BRAF35, and CoREST, which bridges LSD1 to nucleosomes, thereby promoting LSD1-mediated H3K4me removal^29,30,32–34^. The PHF21A-LSD1 complex is recruited by a TF, REST/NRSF, and suppresses neuron-specific genes in non-neuronal cells^29^. The roles of PHF21A in neurons remain unknown.

Six possible alternative splicing isoforms of PHF21A, termed BHC80-1 to -6, have been reported in mice, yet their functional differences remain largely unknown^35^. The above in silico analysis indicated that BHC80-4 is a neuron-specific isoform, whereas BHC80-6 is the canonical form; we refer to them as PHF21A-n and PHF21A-c, respectively. The PHF21A neuronal microexon generates shorter protein segments than PHF21A-c immediately upstream of the PHD finger, which recognizes H3K4me0 (Figure 1A). Within the brain, the PHF21A microexon was included only in neurons while absent in any major non-neuronal cells of the mouse brain (Figure 1B)^27^. To test which neuronal subtypes express the PHF21A microexon, we turned to RiboTRAP sequencing dataset profiling ribosome-bound mRNA from four neuronal subtypes: CamKII(+) neocortical pyramidal neurons, Scnn1a(+) spiny stellate and star pyramid layer 4 neurons, and two GABAergic interneuron types marked by somatostatin (SST) and parvalbumin (PV)^36^. The PHF21A microexon was expressed in all neuron subtypes at a comparable level (Figure 1C), while the non-neuronal exon was not expressed. RT-PCR analysis indicated that the onset of microexon inclusion in PHF21A and LSD1 was approximately E13.5, coinciding with bulk neurogenesis within the mouse brain (Figure 1D).

**Figure 1:**
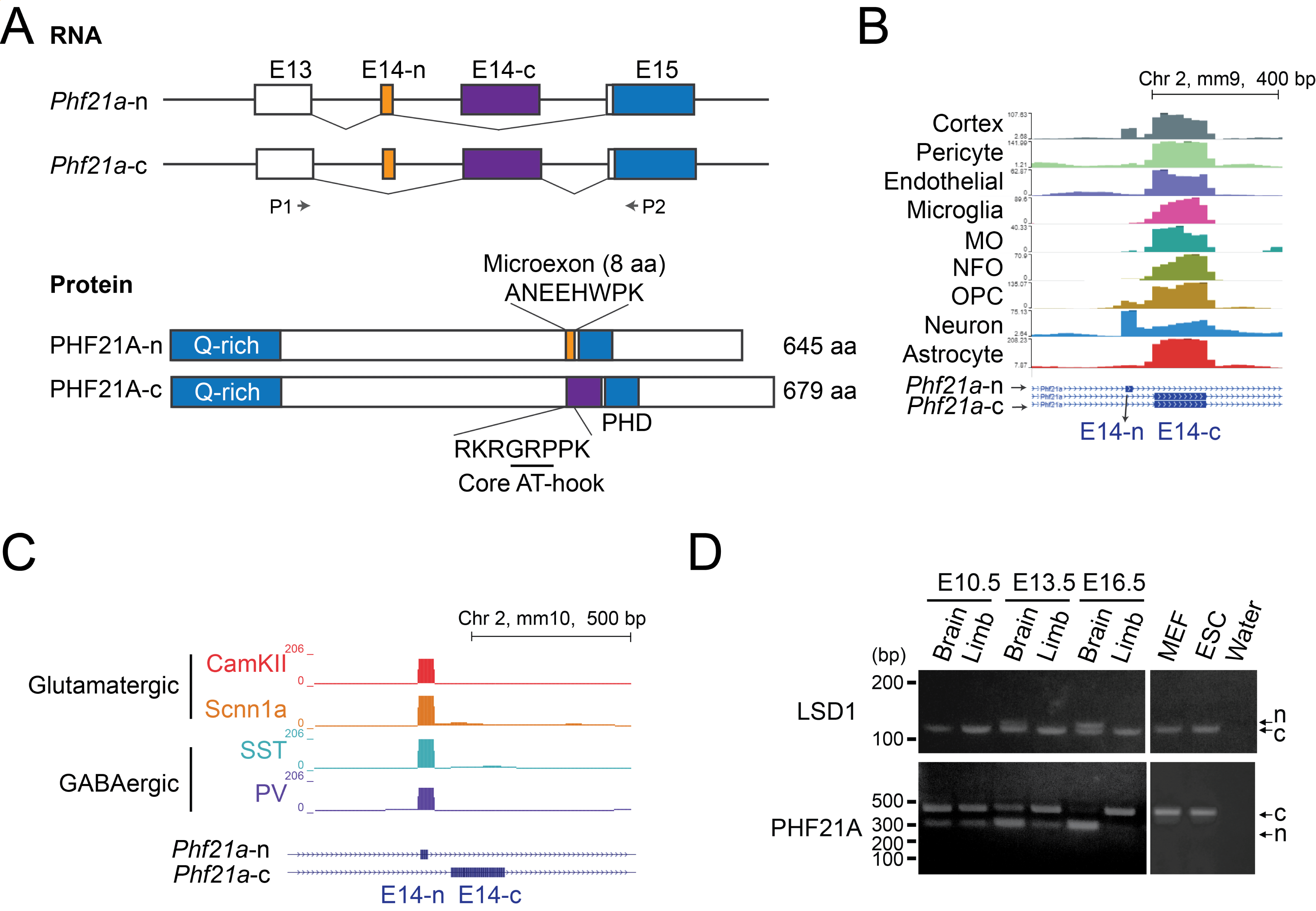
Identification of PHF21A-n. (A) Splicing pattern of *PHF21A* mRNA isoforms and domain organization of canonical and neuronal PHF21A (PHF21A-c, PHF21A-n) proteins. The PHF21A-n neuronal microexon disrupts an AT Hook domain upstream of Plant Homeodomain (PHD) finger. P1 and P2 indicate primers used for the semi-quantitative PCR in panel D. Human and mouse PHF21A share identical splicing patterns, and amino acid sequences of alternative exons. (B) A genome browser view of *Phf21a* splicing events in the mouse cell-type-sorted mRNA-Seq data^27^. (C) PHF21A isoform expression in the four mouse neuronal subtypes. RiboTrap mRNAseq data from the Furlanis et al. paper^36^ were reanalyzed. (D) RT-PCR for *Lsd1* and *Phf21a* alternative isoforms. Both genes show the specification of the neuronal microexon as soon as E13.5 mouse embryonic brains.

In humans, broad brain regions but no other adult tissues express PHF21A-n (Figure 2A). The amino acid sequence of PHF21A-n is 100% conserved between mice and humans (not shown). To examine the developmental time course of human PHF21A-n expression, we differentiated human induced pluripotent cells (iPSCs) into cortical neurons^37–39^ and measured PHF21A-c and PHF21A-n mRNA levels by RT-qPCR (Figure 2B & 2C, Figure S2A). While total PHF21A levels increase continuously during the differentiation process, the ratio of PHF21A-n/PHF21A-c showed a marked increase after neural progenitor cells (NPCs) mature into postmitotic neurons (DIV5 to DIV 20). Still, a substantial fraction of PHF21A adopts canonical form even during neuronal maturation. In this differentiation protocol, we detected significant increases in both neuronal (*DCX*, *NeuN*, *MAP2*) and astrocyte (*GFAP*) marker genes (Figure S2B). To test the presence of PHF21A-c is due to glial cells or the coexpression of canonical and neuronal forms in neurons, we differentiated iPSCs into a relatively homogeneous population of glutamatergic and cholinergic neurons with inducible expression of Neurogenin1 and 2 (NGN1/2)^40^. In this system, PHF21A-n became the only detectable form at differentiation DIV4 (Figure 2D & 2E, Figure S2C), confirming mutually exclusive expression of PHF21A-n in neurons and PHF21A-c in non-neuronal cells such as astrocytes. The molecular weight of PHF21A-n and -c proteins matched with those of recombinant proteins expressed in SH-SY5Y cells (Figure 2F). Together, these results demonstrate that PHF21A-n is the dominant isoform in neurons in mice and humans.

**Figure 2:**
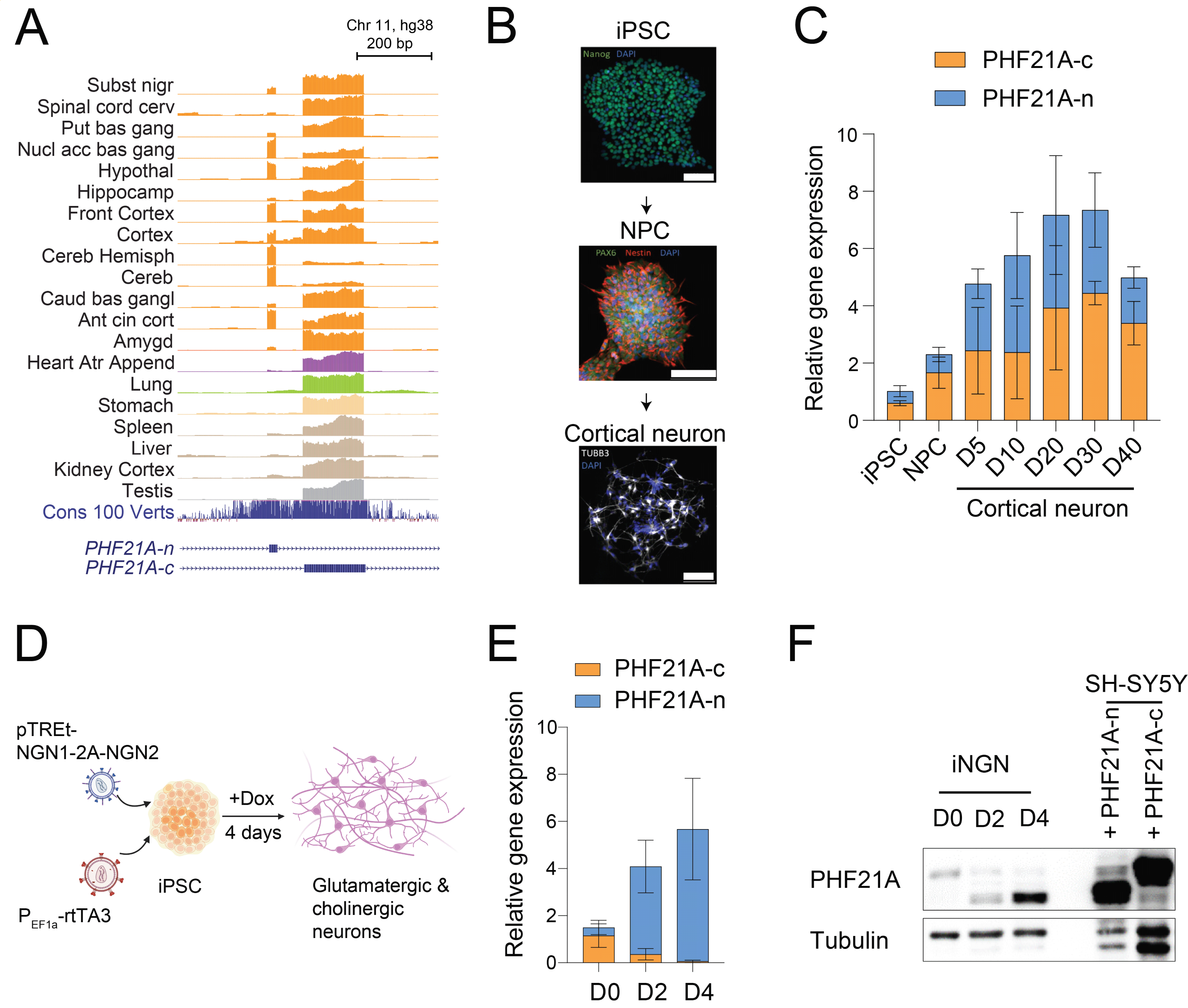
Expression of human PHF21A-n. (A) Brain-specific expression of PHF21A-n in humans based on the GTEx RNAseq data^28^. (B) Schematic of the cell differentiation model. Neural progenitor cells (NPC) and cortical neurons were sequentially generated from induced pluripotent stem cells (iPSC) following established protocols^37–39^. Images are immunofluorescence staining of cells at each stage for some specific markers: Nanog for iPSC, Pax6 and Nestin for NPC, and TUBB3 for neurons. Scale bar: 100 *μ*m. (C) The ratio of PHF21A-c and PHF21A-n mRNA expression levels during neuronal differentiation (n = 3-5, mean ± SEM). (D) Schematic of the engineered iNGN model which relies on the inducible overexpression of NGN1 and NGN2 to rapidly generate a pure population of neurons from iPSC^40^. (E) The ratio of PHF21A-c and PHF21A-n mRNA expression levels at day 0 (D0) (IPSC), day 2 (D2) (precursor stage), and day 4 (D4) (neurons) of iNGN differentiation (n = 7, mean ± SEM). (F) Protein levels of PHF1A proteins during iNGN cell differentiation were analyzed by western blot with a PHF21A-specific antibody. Samples of SH-SY5Y cells transfected with either PHF21-c or PHF21A-n were loaded as control. Tubulin was used as a loading control.

### PHF21A-n splicing disrupts AT-hook-mediated DNA binding

At a protein level, PHF21A-c (∼94 kDa) and PHF21A-n (∼83 kDa) were clearly segregated between the primary neuron and astrocyte cultures (Figure 3A). During the course of this study, another group reported that PHF21A-n is expressed in neuroendocrine prostate cancer and promotes cancer progression^41^. In this study, the alternative segment of PHF21A-c was identified as a putative nuclear localization signal, which is absent in PHF21A-n. As a result, PHF21A-n was localized in the cytoplasm of the cancer cell line. However, our immunofluorescence assays demonstrated that both endogenous and recombinant PHF21A-n, like PHF21A-c, localized at the nucleus in neurons (Fig. 3B & 3C). Furthermore, similar to PHF21A-c in MEF, PHF21A-n in neurons bound to the known interaction partners, including LSD1, CoREST, and HDAC2 (Fig. 3D). To investigate the impact of the neuronal microexon inclusion on the H3K4me0 recognition, we bacterially expressed and purified GST-fused PHF21A-PHD with or without the upstream alternatively spliced region (Figure 3E) and performed histone-peptide binding assays. As expected, PHF21A-PHD without the upstream segment bound to the H3K4me0 peptide, and the interaction was blocked by K4 dimethylation (Figure 3F, lane 4). Despite the alternative sequences nearby the PHD, both PHF21A-c and PHF21A-n showed specific binding to H3K4me0 but not H3K4me2, indicating that two isoforms share the same methyl-histone recognition property (Figure 3F, lanes 5-6).

**Figure 3:**
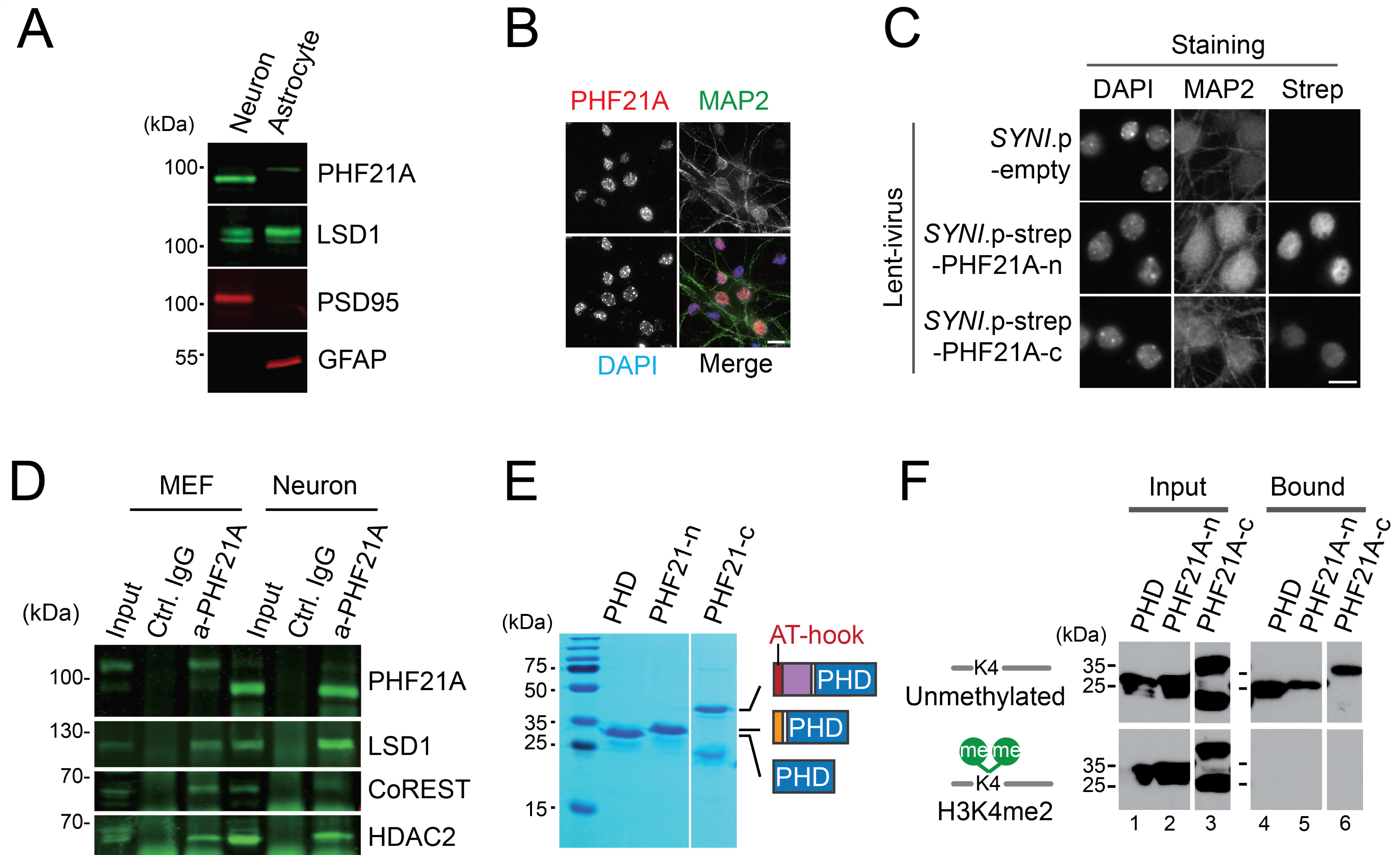
PHF21A-n is a nuclear protein that recognizes H3K4me0, similar to PHF21A-c. (A) Western blot with a primary cortical neuron (DIV10) and astrocyte (DIV21) culture. Anti-PHF21A recognizes N-terminus common to the two isoforms. PHF21A isoforms separate into Tuj-(+) neurons and GFAP(+)-astrocytes. (B) Immunofluorescence of endogenous PHF21A in mouse cortical neurons. (C) Immunofluorescence of recombinant PHF21A. Strep-tagged PHF21A isoforms were expressed in mouse cortical neurons (DIV10) under the hSYNI promoter and detected by the indicated antibodies. Scale bar: 10 *μ*m. (D) Co-IP assays to test the interaction of PHF21A and known partners. Nuclear extracts from MEF and cortical neurons were used. (E) GST-PHF21A-PHD/AT-hook fragments were expressed in E. coli and affinity purified. (F) Histone peptide pull-down assays. Biotinylated histone H3 peptides (a.a. 1-20) with or without H3K4me2 were incubated with GST-PHF21A fragments, pulled down by streptavidin affinity resin. Bound proteins were detected by Western blotting with an anti-GST antibody.

We then realized that the alternative PHF21A-c contained an uncharacterized AT-hook motif, which could potentially bind to the minor groove of AT-rich DNA^42^, and the neuronal microexon disrupts this AT-hook (Figure 1A and B). We, therefore, tested the DNA binding function of PHF21A isoforms through Electrophoretic Mobility Shift Assays (EMSA). We found only PHF21A-c but not PHF21A-n was able to bind to Widom601 DNA (Figure 4A) or the Widom601 recombinant nucleosome (Figure 4B). The slight shift of nucleosome by higher protein concentrations can be attributed to the PHD-H3 tail interaction (Figure 4B, lanes 6 & 11). We also found robust DNA binding by PHF21A-c but not by PHF21A-n when we used nucleosomes from 293T cells or DNA thereof, suggesting that the DNA binding by PHF21A-c is independent of nucleotide sequences at a higher concentration (Figure 4C and D).

**Figure 4.**
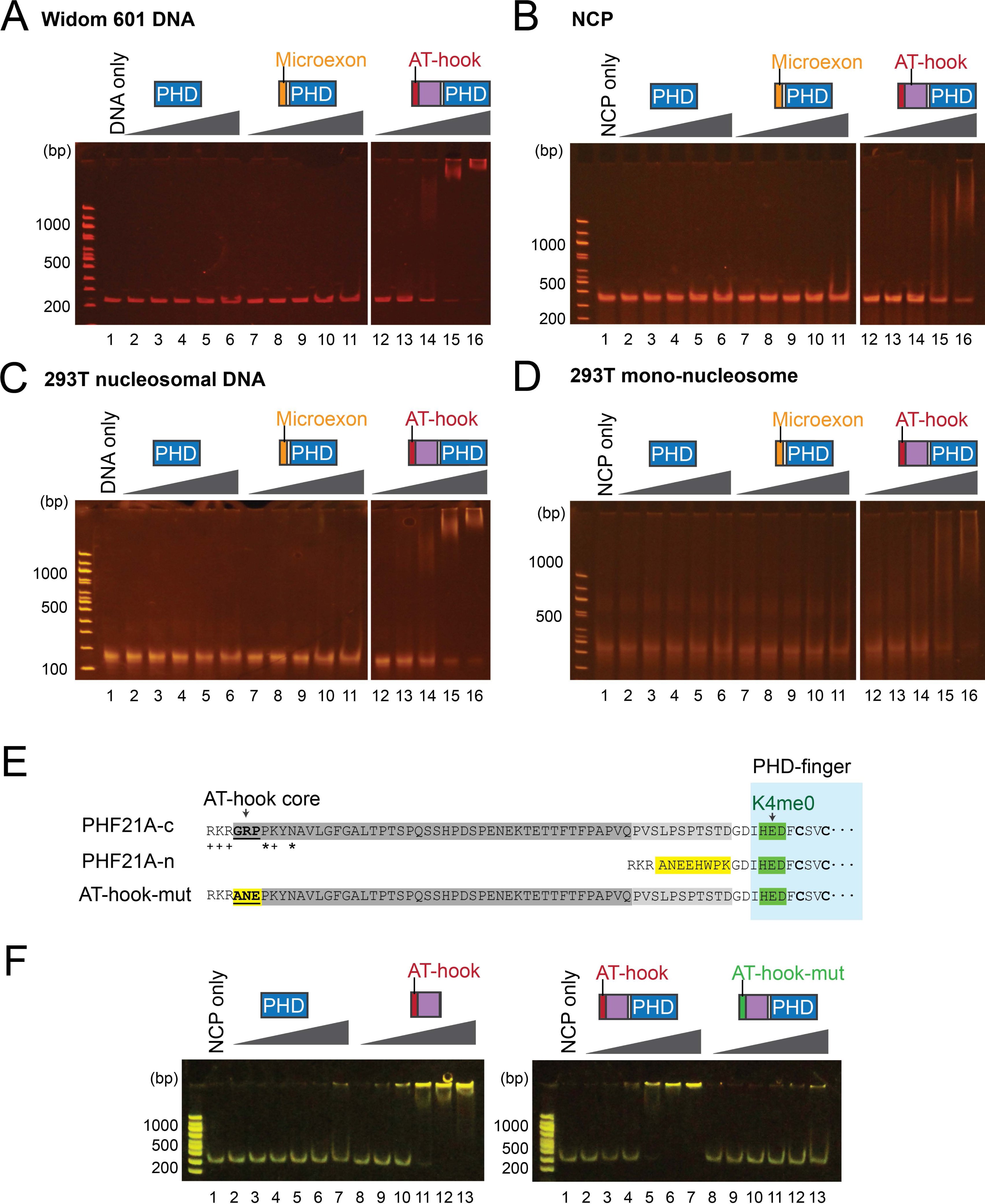
PHF21A-c is a DNA-binding protein, and neuronal splicing ablates the DNA-binding function. (A) EMSA with Widom 601 DNA and PHF21A fragments. (B) EMSA with recombinant nucleosome core particles (NCP). (C) EMSA assay using nucleosomal DNA extracted from 293T-derived mono nucleosomes. (D) EMSA assay using 293T-derived mononucleosomes. (E) The amino acid sequence of PHF21A isoforms and the AT-hook-mutant. The core AT-hook motif GRP was replaced with ANE, the corresponding PHF21A-n sequence. Basic (+) and polar (*) amino acids that potentially contribute to DNA binding are denoted (F) NCP-EMSA with PHD only, AT-hook only, AT-hook-PHD, and the AT-hook-mutant.

To evaluate the roles of the AT-hook within the alternative PHF21A-c segment, we mutated three core residues of the AT-hook (GRP) to those of PHF21A-n (ANE) (Fig. 4E). The three amino-acid substitutions largely abolished the nucleosome binding (Fig. 4F lanes 8-13). Additionally, AT-hook alone had a much stronger affinity to Widom 601 nucleosome than the PHD finger (Fig. 4F). These data highlight a new role for AT-hook-mediated DNA binding by PHF21A, providing a dominant affinity for engaging nucleosomes, and demonstrate that the neuronal microexon selectively abolishes the DNA-binding function leaving PHD-driven H3K4me0 recognition intact.

### Impact of neuronal exons on LSD1-CoREST-PHF21A complexes

To investigate the biochemical properties of the canonical and neuronal PHF21A and LSD1 as a complex, we reconstituted complexes consisting of LSD1, CoREST, and PHF21A, prepared with the insect cells transduced with baculovirus system. The reconstituted complexes are either bipartite, consisting of LSD1 and CoREST (LC), or tripartite, consisting of PHF21A, LSD1, and CoREST (PLC) (Figure 5). Canonical complexes (LC-c, PLC-c) included LSD1-c and PHF21A-c, whereas the neuronal complexes (LC-n, PLC-n) contained LSD1-n and PHF21A-n. We first tested whether the neuronal components can assemble into a functional complex. To this end, LSD1 was MBP-tagged for the bipartite LC complexes. We then co-expressed the components in Hi5 cells, affinity-purified the complex with MBP resin, and subjected the purified complexes to the size exclusion column. Stoichiometric complexes were eluted with either canonical or neuronal components, indicating that LSD1-n and PHF21A-n can assemble into a complex akin to the canonical counterpart (Figure 5B, lanes 1-5). The efficient assembly of neuronal complexes is consistent with the Co-IP results (Figure 3D).

**Figure 5.**
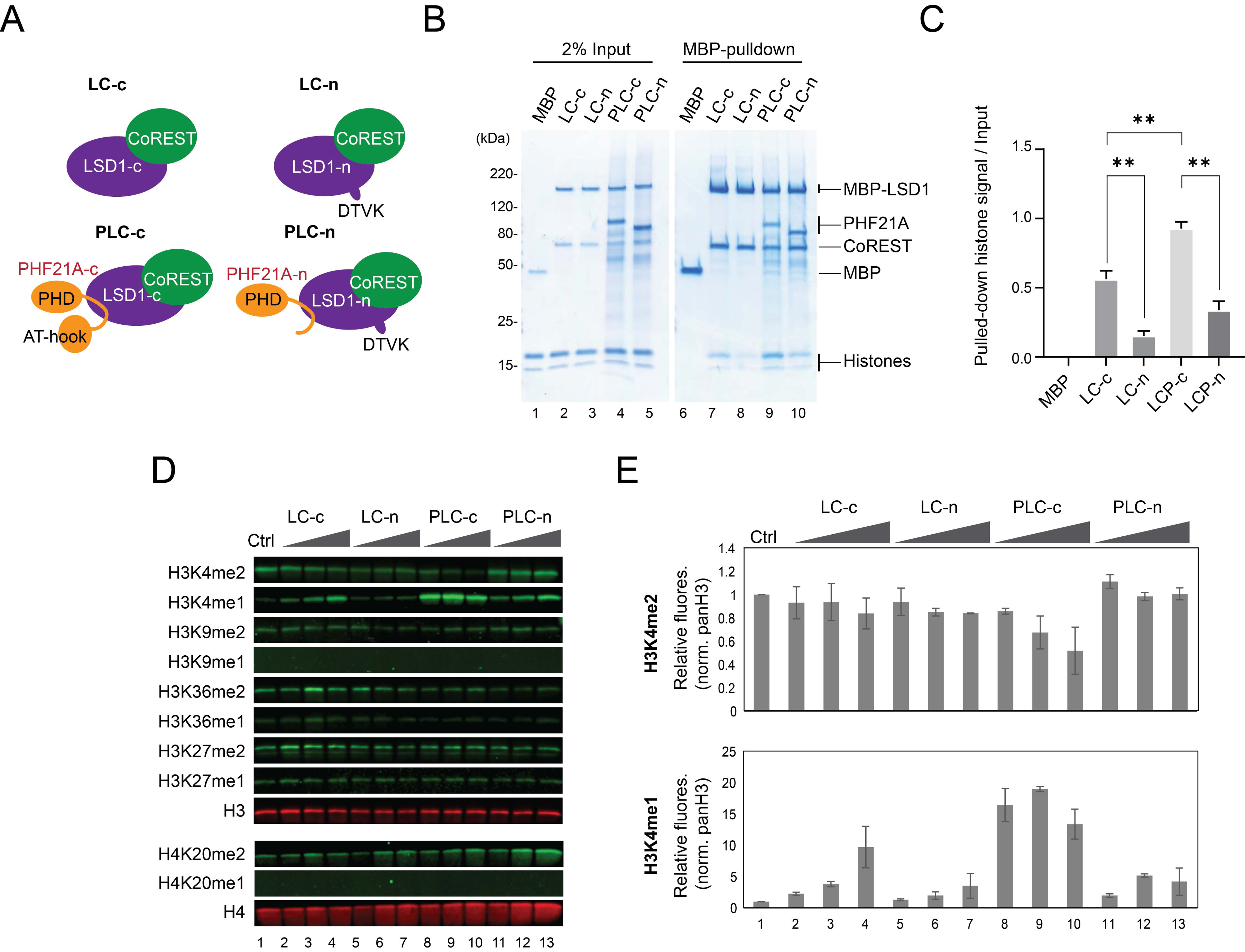
Neuronal LSD1-PHF21A complexes are hypomorphic H3K4 demethylation machinery in vitro. (A) Schematics of reconstituted complexes using protein prepared with Sf21 cells. (B) Input (lanes 1-5), LSD1 and CoREST were co-expressed in Sf21 cells where LSD1 N-terminus was fused to the MBP tag, pulled down by amylose resins, and further purified by a size-exclusion column. The bipartite LSD1-CoREST complex was then incubated with unmodified recombinant mononucleosomes (NCP) with H3K4M substitution with or without recombinant PHF21A, pulled down with amylose resins, separated by SDS-PAGE, and visualized with coomassie staining. (C) Quantification of pulled-down NCP. Pulled-down histones’ band intensity was normalized with the input histone signals (n=3, mean ± SEM). P values were calculated using one-way ANOVA followed by the Tukey test with multiple comparison corrections. *P<0.002, **P<0.0001 (D) Demethylation assays. The reconstituted complexes were incubated with the designer nucleosomes carrying the denoted histone di-methylation and analyzed by quantitative Western blot using the antibodies against corresponding methylations. (E) Quantification of the Western blots. Signals were normalized to H3 levels measured by anti-pan H3 or H4 antibodies (n = 3 independent reactions, mean ± SEM).

We then tested the affinity of the complexes to the nucleosome. The reconstituted complexes were incubated with recombinant Widom 601 nucleosomes, carrying an H3K4M mutation to mimic the methylated state^43^, pulled down by the MBP affinity resin. The PLC-c complexes bound to the nucleosome more strongly than the LC-c complex, indicating that PHF21A stabilizes the LSD1 onto the nucleosome (Figure 5B, lanes 7 vs. 9, and 5C). Interestingly, the nucleosome binding of LC-n was reproducibly weaker than that of LC-c, suggesting that neuronal splicing dampens affinity between the LSD1-CoREST complex and the nucleosome (Figure 5B, lanes 7 vs. 8, and 5C). As tripartite complexes, PLC-n again displayed significantly weaker nucleosome interactions than PLC-c (Figure 5B, lanes 9 vs. 10, and 5C). The results indicate that the microexons of PHF21A and LSD1 concertedly dampen the affinity of the LSD1-CoREST-PHF21A complex to the nucleosome.

Next, we examined the enzymatic activity of the reconstituted complexes using recombinant designer nucleosomes that uniformly carried di-methylation of the major five histone lysines: H3K4me2, H3K9me2, H3K27me2, H3K36me2, or H4K20me2. Quantitative fluorescent Western blots were used to measure methylation levels of the nucleosome after reactions. The PLC-c complex showed significantly stronger H3K4me2 demethylase activity than the LC-c complex, indicating that PHF21A facilitated the reaction (Figures 5D & E). Notably, PLC-c at lower concentration produced a high H3K4me1 level, which decreased at a higher concentration of PLC-c, indicating a successive H3K4me2 and me1 removal (Figure 5D & E, lanes 8-10). While LC-n mediated H3K4 demethylation was barely detectable, PLC-n did exhibit a weak activity judged by the production of H3K4me1 and reduction of H3K4me2 (Figure 4C & 4D, lanes 11-13). Demethylation of H3K9me2, H3K27me2, or H4K20me2 was not observed in any of the reactions (Figure 5D). Taken together, the reconstituted neuronal complex PLC-n is identified as a hypomorphic H3K4 demethylase complex.

### A human *PHF21A* mutation in syndromic autism

*PHF21A* haploinsufficiency is responsible for intellectual disability and craniofacial abnormalities seen in Potocki-Shaffer syndrome (MIM: 601224)^44–47^. PHF21A is an autism-associated gene^48^. However, since most of the reported genetic lesions alter both PHF21A-c and PHF21A-n, the specific contribution of PHF21A-n to pathogenesis remains unknown. A recently-described single *de novo* substitution *PHF21A* c.1285G>A in a syndromic autism patient^45^ (Figure 6A) provides an opportunity to probe the role of PHF21A-n in human brain development. This mutation occurs at the last nucleotide of exon 13 directly upstream of the neuronal microexon, therefore, leading to four possible molecular consequences: a splicing defect of PHF21A-c or PHF21A-n, and two distinct amino acid substitutions in the two isoforms, i.e. p.Gly429Ser in PHF21A-c and p.Ala429Thr in PHF21A-n (Figure 6A). Since the mutated PHF21A-c-Gly429 is one of the three AT-hook core amino acid residues, the substitution was suspected of interfering with the DNA-binding function^45^.

**Figure 6.**
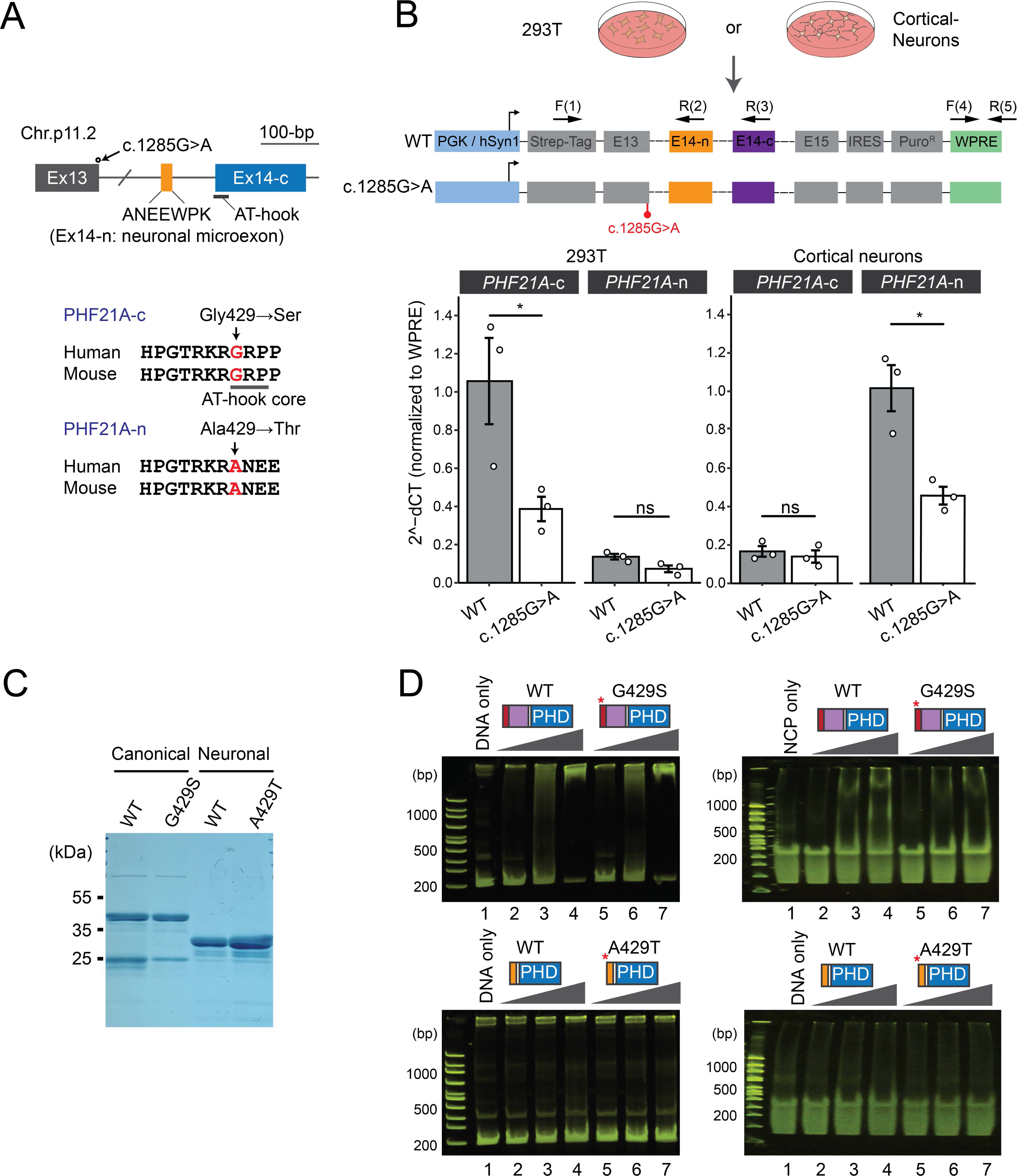
The consequences of PHF21A c.1285G>A mutation of a syndromic autism patient. (A) The schematics of c.1285G>A substitution and distinct amino acid substitutions between the c and n forms. (B) mini-gene splicing assay in 293T cells. Upper panel: Schematics of the mini-gene-splicing assay. Expression of the PHF21A mini-genes wase driven by the hPGK promoter in 293T cells and the hSYN1 promoter in cortical neurons and quantified by RT-qPCR with the primers specified. “E” denotes exons. Bottom panels: qPCR quantification of spliced exons in 293T cells (left) or cortical neurons (Right panel). The abundance of spliced exons was normalized to the expression of the WPRE sequence (n = 3, Mean ± s.e.m., * P<0.05, ns: not significant, two-tailed student t-test with equal variance). (C) WT and mutant PHF21A-PHD fingers expressed and purified from E.coli. (D) EMSA assay using Widom601 DNA (left panels) or NCP (right panels) with canonical WT and the mutant proteins.

We, therefore, set out to test these possibilities. First, we examined the impact of the *PHF21A* mutation on splicing by introducing WT and the mutant mini *PHF21A* gene carrying the c.1285G>A to 293T cells or mouse cortical neuron culture (Figure 6B). The PGK and SynI promoters were used for expressing the mini-genes in non-neuronal and neuronal cells, respectively. The splicing products were quantified by RT-qPCR using primers specific to the recombinant mini-gene sequences. PHF21A-c was the dominant splicing product in 293T cells, while PHF21A-n was dominant in cortical neurons, indicating that the assay recapitulates the cell-type specific splicing events. The mutation led to a 63 % and 55 % reduction of canonical and neuronal splicing products (Figure 6B). The result indicates that mutant protein can be expressed in the cell albeit at a lower level. Therefore, we next examined the impact of the amino acid substitutions on DNA binding. The mutant proteins were expressed at a comparable level with WT, suggesting that the protein stability was not perturbed by the mutations (Figure 6C). Though the AT-hook core “GRP” is responsible for the DNA-binding function of PHF21A (Figure 3B & 3C), we did not find noticeable differences between WT and Gly429Ser mutant in either DNA or nucleosome binding (Figure 6D). The Ala429Thr substitution on neuronal isoform did not change the lack of DNA-binding capability of PHF21A-n. These results suggest that PHF21A c.1285G>A mutation interferes with the production of both PHF21A-c and PHF21A-n and that the one amino acid change in AT-hook is not sufficient to alter DNA binding function. Thus, a reduced PHF21A-n level is likely a contributing factor to the neurodevelopmental traits of the affected individual.

## Discussion

In this study, we report neuron-specific chromatin dynamics, in which two coordinated alternative splicing events within the LSD1-PHF21A complex alter the intrinsic activity of the complex. Prior studies have described unique chromatin regulation in neurons. For example, CpA DNA methylation, catalyzed by DNMT3A, finetunes gene expression for neuronal maturation^49^. In addition, the nucleosome spacing is ∼17-bp shorter in neurons than that of astrocytes^50^. As discussed earlier, neuron-specific chromatin-regulatory complexes have been described for BAF and TFIID complexes. The two complexes carry unique components in different cell types, such as BAF53b and TAF9B in neurons^7,11^. These unique subunits are only present, or more abundant, in neurons or during specific developmental time points. Mechanistically, these unique subunits possibly tether the core enzymatic components, such as ATPases, to specific genomic loci, thereby imparting specificity of gene regulation^10,51^. This mechanism, therefore, can be viewed as an extension of the TF-mediated target specificity. In contrast, the data presented in this study demonstrate that cell-type-specific complexes can have altered intrinsic function.

The roles of the LSD1-PHF21A complex have been debated on two fronts. The first issue is the substrate specificity of LSD1-n: H3K4me^17^, H3K9me^20^, or H4K20me^24^. The three reports used different methods to measure the activity of LSD1-n. Zibetti et al. used bipartite complexes of LSD1 and CoREST C-terminus and synthetic H3 peptides. Laurent et al. isolated LSD1-n-containing complexes from differentiating SH-SY5Y cells and measured their activity towards H3 peptides or calf histones. Wang et al. used bacterially-expressed LSD1-n or LSD1-n-CoREST mixtures and measured their enzymatic activities on H3 peptides, histones, and cellular nucleosomes. The present work used tripartite complexes consisting of LSD1, CoREST, and PHF21A expressed in insect cells and designer recombinant nucleosomes as substrate. Designer di-methyl nucleosomes used in this study provide a higher sensitivity over the cellular nucleosomes because abundant H3K4me1 present in cellular nucleosomes can mask demethylation-derived H3K4me1. In addition, demethylation-mediated reduction of preexisting H3K4me1 and newly generated H3K4me1 offset each other. Therefore, our results provide reliable evidence that the neuronal tripartite complex, consisting of LSD1-n, PHF21A-n, and CoREST, is incapable of demethylating H3K9me or H4K20me but retains weaker activity towards H3K4me than the canonical counterpart (Figure 5). However, the current study does not exclude the possibility that the neuronal complex might have a different substrate specificity in vivo by interacting with additional factors, as reported by Laurent et al.

The present study provides insights into the molecular etiology of neurodevelopmental disorders. Beyond the specific syndromes associated with PHF21A and LSD1 mutations, the loss of function in chromatin regulators has emerged as a major genetic basis of autism spectrum disorder and intellectual disability^55^. It is tempting to speculate that neuron-specific chromatin dynamics contribute to the brain’s vulnerability to chromatin deregulation. Our investigation on PHF21A 1285G>A substitution provides support for an important role of PHF21A-n in human brain development. Though the negligible impact of Gly429Ser on DNA binding was inconsistent with the previous prediction by Kim et al.^45^, known mechanisms of DNA recognition by HMGA AT-hook can explain the lack of impact. The corresponding glycine in HMGA AT-hook provides hydrophobic contact to the DNA base and ribose backbone in a solution structure^56^ or a hydrogen bond to thymine in a crystal structure^57^. The Gly429Ser substitution was predicted to interfere with DNA binding based on the hydroxyl group of serine, creating a repulsive negative charge to the DNA phosphate^45^. Meanwhile, the HMGA AT-hook core, PRGRP, has only a weak affinity to DNA with a micromolar range. Basic and polar amino acids, neighboring the core, contact the DNA phosphate backbone, thereby achieving nanomolar affinity^42,56,58^. Such basic and polar amino acids are present adjacent to the PHF21A AT-hook core motif (Figure 4E). Thus, the additional DNA contacts outside the core motif might make Gly429Ser substitution tolerated. In addition, the glycine-serine substitution, keeping the small amino acid size, unlikely disrupts the embedding of the AT-hook core motif to the DNA minor groove.

Prior mechanistic studies of microexon splicing point to possible molecules controlling PHF21A-n splicing. The majority of microexon splicing relies on nSR100, a neuron-specific splicing factor^12,59^. Indeed, PHF21A microexon is present in the list of nSR100-dependent neuronal exons^59^. While nSR100 promotes the inclusion of neuronal microexons, polypyrimidine tract binding protein PTB suppresses neuronal splicing events. PTB knockdown leads to the altered splicing patterns of PHF21A in HeLa and N2A cells^59^. In addition to PTB, LSD1-n splicing is also regulated by nSR100 and NOVA^21^. The UGC sequence at the 3’-splice site is an nSR100 binding motif^12,60^, and we observed that the upstream of PHF21A-n exon has several UGCs. Thus, PHF21A-n splicing is likely coordinated by nSR100 and PTB.

The ASD-associated PHF21A mutation c.1285G>A offers an insight into the role of PHF21A-n in human brain development. Mechanistically, the mutation decreases expression of both PHF21A-c and PHF21A-n (Figure 6), suggesting a binding impairment of general spliceosome components such as the U1 snRNP as opposed to impairment of the above neuronal splicing factors. The patient with this mutation exhibits both neurological phenotypes, such as autism and intellectual disability, along with non-neuronal phenotypes, including clinodactyly^45^; therefore, a reduced dose of both PHF21A-c and PHF21A-n likely contributes to the disease phenotypes. Meanwhile, the little effect of PHF21A-c Gly429Ser substitution on DNA binding reveals that functional interference of PHF21A-c may not contribute to the cognitive deficits of the affected individual. However, why PHF21A needs to be a neuronal form instead of the canonical form in neurons remains an open question. A swap mutant in which PHF21A-n is replaced with PHF21A-c in neurons would be useful to address this problem in the future.

Our in silico analyses indicated that neuron-specific splicing events occur in more than a dozen of chromatin regulators (Table 1). Thus, LSD1-PHF21A-complex is unlikely unique in having neuron-specific intrinsic function; instead, splicing-mediated modulation of chromatin-regulatory activity represents a general mechanism by which neurons generate their unique chromatin landscapes. Our data demonstrate that PHF21A neuronal splicing is strictly conserved between mice and humans (Figures 1 and 2). Intriguingly, some neuronal splicing events are species-specific, such as G9a only being found in mice, raising a possibility that neuronally-spliced chromatin factors might contribute to the diversity of brain development and functions across species. Thus, our work opens a new avenue of research to explore neuron-specific chromatin regulatory activities for their roles, evolution, and human disease relevance.

## Acknowledgment

We thank the Iwase laboratory members for their helpful discussions and critical review of the data. We also thank Isabel Wellik for her experimental help with the EMSA assays of the PHF21A mutant. This work was funded by an NIH National Research Service Award T32-HD079342 (University of Michigan Predoctoral Career Training in the Reproductive Sciences Program) from the National Institute of Child Health and Human Development (NICHD) (to RSP and KMB), an NIH National Research Service Award F31NS103377 (to RSP), University of Michigan Medical Scientist Training Program Fellowship (T32 GM007863 to RSP), Rackham Graduate School Pre-doctoral Research Grant (to RSP and KMB), an NIH Award R01NS089896, R01NS116008, R21NS104774, R21NS125449, and R21MH127485 (to SI).

## Author Contributions

RSP and SI initiated the project. RSP, MN, SA, MCG, BZ, OD, JMM, JSG, YM, and LB carried out experiments and interpreted and collected data. KMB performed RNA-seq analyses. USC supervised in vitro reconstitution experiments. BL supervised human neuron differentiation studies. RSP and SI wrote the manuscript draft, and all authors edited it as needed and approved it. SI supervised the project and acquired funding.

## Declaration of Interests

The authors declare no conflict of interest.

## Supplementary Figure Legends

**Figure S1.**
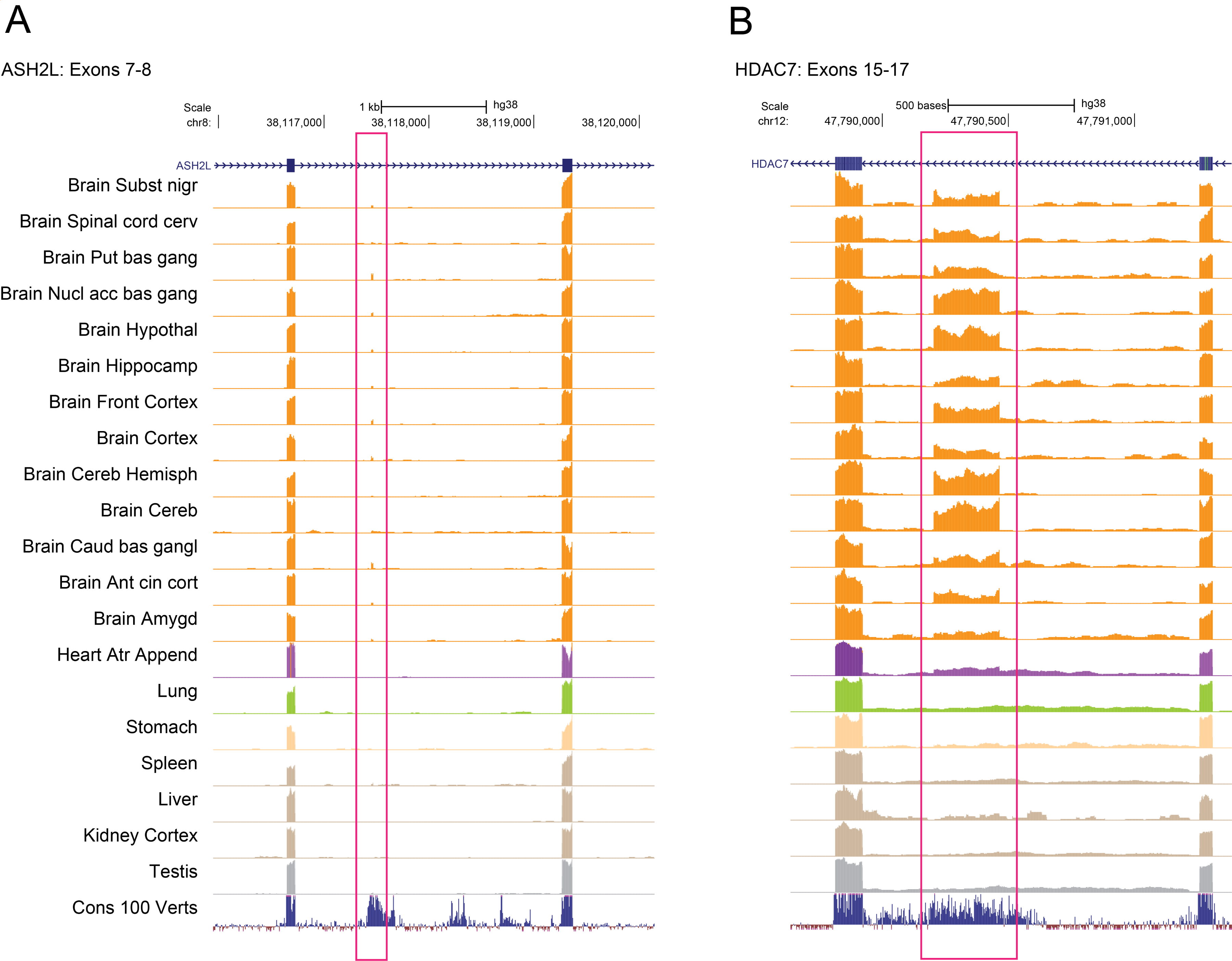
Evidence of unannotated neuronal splicing events in ASH2L and HDAC7. Read coverage of Human GTEx RNAseq data^28^ is displayed in the UCSC genome browser for ASH2L (A) and HDAC7 (B). The two genes undergo splicing events unique to brain tissues, as highlighted in pink rectangles.

**Figure S2.**
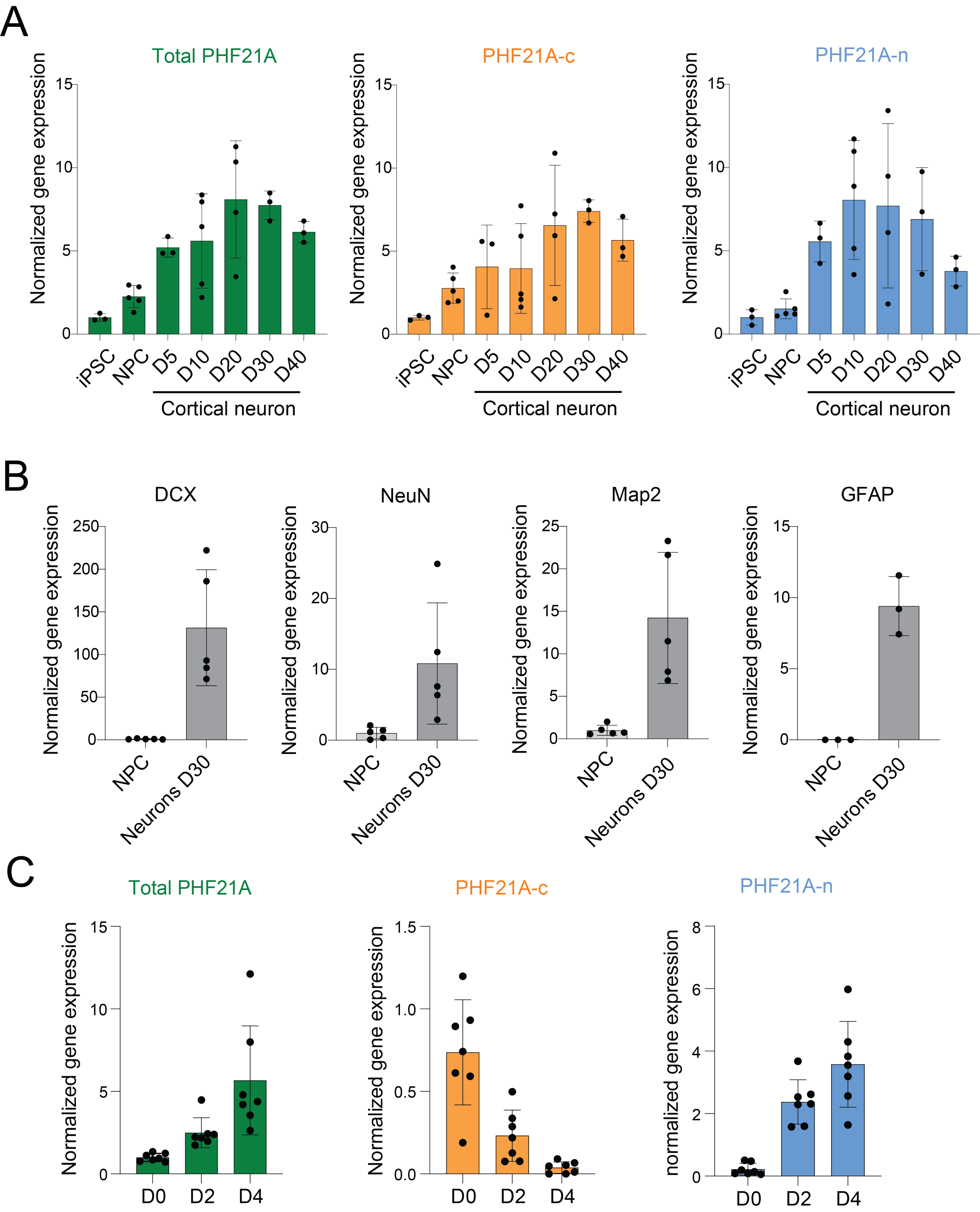
Expression of human PHF21A-n. (A) mRNA expression levels of total PHF21A, PHF21A-c, and PHF21A-n were measured by quantitative PCR (qPCR) at different time points during the sequential differentiation of iPSC into NPC and neurons presented in Fig. 2B. (B) mRNA expression levels of *DCX*, *NeuN*, *Map2*, and *GFAP* were measured by qPCR in NPC and neurons (day 30 of differentiation). (C) mRNA expression levels of total PHF21A, PHF21A-c and PHF21A-n were measured by qPCR at D0 (iPSC), D2 (precursor stage) and at D4 (neuron) of iNGN differentiation.

## Materials and Methods

### Identification of neuron-specific splicing events in chromatin regulators

Chromatin regulators, as defined by EpiFactors^25^ were intersected with neuron-specific splicing patterns as defined previously^12^, which led to 115 chromatin regulators that undergo neuron-specific splicing. We then filtered the splicing events that did not satisfy manual inspection of read coverage data using the UCSC genome browser. For this filtering step, we used both mouse and human GTEx data^27,28^, and obtained the 14 chromatin regulators genes. Those splicing events that fall into a known protein domain defined by the NCBI Conserved Domain Database^61^ were depicted through their domain structures. We assessed alternative splicing in different neuronal cell types using reporter-driven RiboTRAP RNA sequecing^36^. We aligned sequencing reads to the mm10 genome with STAR (v2.6.1a), generated bigWigs using bamCoverage (v3.2.0), and visualized the data using the UCSC genome browser.

### Constructs and lentiviral vectors

Mammalian expression vectors for PHF21A and LSD1 were cloned first into a pENTR vector by Gibson assembly. Subsequently, these entry plasmids were recombined into mammalian destination plasmids using Gateway LR Recombination (Thermofisher) into pcDNA3.1 (Addgene #52535) and pLIX-402 (Addgene #55169). For bacterial protein expression, cDNA constructs were cloned into pGEX-4T1 (Addgene #2876). For insect cell protein expression, cDNA constructs were cloned into a pFastBac Dual Expression Vector (Thermo Fisher) using Ligation Independent Cloning^62^. AT-hook mutations were introduced into the above plasmids using site-directed mutagenesis. All plasmids are available upon request.

### Cell culture

#### Primary Neurons

The primary neuron culture was performed as previously described^63^. Timed pregnant female mice were sacrificed at day E16.5, and embryonic brains were harvested. The cortices and hippocampi were microdissected, treated with 2.5% Trypsin (Invitrogen, 15090), quenched by FBS, and then treated with 1% DNaseI (Sigma, DN-25) to dissociate brain tissue into neurons. Cells were counted and then plated in Neurobasal Media (Gibco, 21103049) with 1x B27 (Gibco, 17504-044), 0.5 mM GlutaMax (Gibco, 21985), 25 *μ*M beta-mercaptoethanol, 1x Penicillin-Streptomycin. Before plating, plates were coated with Poly-D-lysine hydrobromide (PDL, Sigma P7886, mw 30,000-70,000) overnight at 37°C and then washed with water three times. Neuron culture cells were fed every 3-4 days by replacing half of the above media with new media. For molecular analyses, Cytarabine (10 *μ*M, aka AraC, Tocris) was added to the culture at DIV2 or DIV3 to eliminate the non-neuronal cell growth.

#### MEFs

E16.5 mouse carcasses were eviscerated, minced, and incubated with 0.25% trypsin (Gibco 25200072) for 30 minutes. Cells were quenched and then plated in DMEM (Gibco 11995065) with 10% FBS, 1% Penicillin-Streptomycin (Gibco 15140122), and 1x GlutaMax (Gibco 35050061). Cells were grown out for 3-5 days, were passaged once, and then were harvested for most experiments.

#### Human induced pluripotent cell lines (iPSCs)

Human iPSCs used in this study were cultured on Matrigel-coated plates (Corning) using mTeSR1 media (StemCell Technologies) with daily medium change. Once reached 80% confluency, cells were passaged using a 5 min incubation with Accutase (StemCell Technologies) on Matrigel-coated plates in mTeSR1 media supplemented with 10 *μ*M of Y-27632 (Cayman Chemicals). SH-SY5Y cells were cultured in DMEM (Wisent) supplemented with 10% fetal bovine serum (Avantor).

#### Sequential differentiation model

The iPSC line AJG-001-C4 (37 years old, healthy male) was bought from the C-BIG Repository. Induction of iPSC into neural precursor cells (NPC) was performed as previously described with some modifications^37–39^. Approximately 80% confluent iPSC were dissociated into a single-cell suspension using Accutase (StemCell Technologies) and plated on Matrigel-coated plates (Corning) at a density of 30 000 cells/cm^2^ in a neural precursor cell differentiation medium (NPCM) supplemented with 10 *μ*M Y-27632 (Cayman Chemicals) and 0.1 *μ*M compound E (Cayman Chemicals). The medium was changed daily with NPCM +0.1 *μ*M compound E. On day 7, NPCs were passaged 1:4 using Accutase on Matrigel-coated plates in NPCM + Y27632. The medium was changed daily with NPCM thereafter. NPCM is composed of Neurobasal:DMEM/F12 (1:1) supplemented with 1% N2-A (StemCell Technologies), 2% B27 without retinoic acid (Life Technologies), 1% Glutamax (Gibco), 1% Penicillin-Streptomycin (Wisent), 10 ng/ml hLIF (StemCell Technologies), 5 *μ*g/ml BSA (Sigma), 4 *μ*M CHIR99021 (Cayman Chemicals) and 3 *μ*M SB431542 (Cayman Chemicals). NPCs were differentiated into cortical neurons using previously described protocols with some modifications. Approximately 90% confluent NPCs were dissociated into a single-cell suspension using Accutase and plated on 40 *μ*g/ml Poly-L-ornithine (Sigma) and 20 *μ*g/ml laminin-coated (Sigma) plate at a density of 20,000 cells/cm^2^ in the neuronal differentiation (NeuroDiff) medium with 10 *μ*M Y-27632. The medium was changed with NeuroDiff 24h after plating and changed every 2-3 days thereafter. NeuroDiff medium is composed of Neurobasal:DMEM/F12 (1:1) supplemented with 1% N2-A (StemCell Technologies), 2% B27 (Life Technologies), 300 *μ*M dibutyryl-cyclic AMP (StemCell Technologies), 20 ng/ml BDNF (Peprotech), 20 ng/ml GDNF (Peprotech) and 200 nM L-ascorbic acid (Sigma).

#### iNGN model

The iNGN cell line was directly differentiated into neurons by the addition of doxycycline as previously described^40^. Briefly, iNGN cells were dissociated into a single-cell suspension using TrypLE Express Enzyme (1X), no phenol red (Gibco), and plated on Matrigel-coated plates at a density of 30,000 cells/cm^2^ in mTeSR1 media supplemented with 10μM Y-27632 and 100μM Revitacell supplement (Gibco). The medium was changed to mTeSR1 + 1 μg/ml of doxycycline (Sigma) after 24 hrs to initiate the differentiation process. The cells were harvested for RNA and protein collection at three different time points: Stem cell (D0) stage, Progenitor Neuronal cells (D2) exactly 48hrs after induction with doxycycline, and at bipolar neurons (D4), exactly 96 hrs after induction.

### RNA isolation and quantitative PCR

Total RNA was isolated using the Quick-RNA™ Miniprep Kit (Zymo Research) according to the manufacturer’s protocol. The RNA concentration was determined using a Nanodrop system (Thermo Fisher Scientific). Total RNA (1 *μ*g) was used to generate cDNA using the Biorad iscript RT supermix RT-qPCR kit (Biorad) following the manufacturer’s instructions. The expression of different target genes was validated by quantitative PCR (qPCR), using the Azure Cielo 3 system (Azure Biosystems, Dublin, CA, USA). The reactions were performed with the PerfeCta SYBR Green SuperMix (QuantaBio, Beverly, MA, USA) as recommended by the manufacturer. Real-time PCR was performed with a hot start step of 95 °C for 2 min followed by 40 cycles of 95 °C for 10 s, 60 °C for 10 s, and 72 °C for 20 s. Analysis was performed using Azure Cielo software (v1.0.0.285, Azure Biosystems, Dublin, CA, USA). The mouse *Phf21a* and *Lsd1* cDNA were amplified with these primers (Figure 1). *Phf21a*-fwd: ACAACAGCAAACCCGGTGTA, Phf21a-rev: TTTTCAGGGGCGGCTCTAAG, *Lsd1*-fwd: TGCCCACTTTATGAAGCCAA, *Lsd1*-rev: ACTGGTGACTAAGGTAAGAAGTGG. The relative expression of human genes was normalized by that of *GAPDH* and *MRPL19*. The specific human primers used for the amplification are as follows: Total PHF21A F: CAGGTTCACCAGAAACTGGTTGCTCAAATGA, Total PHF21A R: CAGGTTCTT CCGTAGCTGTTCAACTACTC, PHF21A-c F: GACAGAGACCACATTCACTTTCCCTGCA, PHF21A-n F: GAGAGCCAATGAGGAACACTGGCCAAAG, PHF21A-n/C R: CAGTCCAAATGATATACACGGGAACATGTGTC, DCX F: TTGCTGGCTGACCTGACGCGATC, DCX R: TTACGTTGACAGACCAGTTGGGATTGAC, NEUN F: GTCGTGTATCAGGATGGATTTTATGGTG, NEUN R: ATGGTTCCAATGCTGTAGGTCGCC, MAP2 F: GGCGGACGTGTGAAAATTGAGAGTGTAA, MAP2 R: ACGCTGGATCTGCCTGGGGAC, GFAP F: GGCCCGCCACTTGCAGGAGTACCAGG, GFAP R: CTTCTGCTCGGGCCCCTCATGAGACG, MRPL19 F: GGAGAAAAGTACTCCACATTCCAGA, MRPL19 R: TGGGTCAGCTGTAGTAACACGA, GAPDH F: TGGGATTTCCATTGATGACAA, GAPDH R: CCACCCATGGCAAATTCC.

### CoIP and Western blot

Immunoprecipitation followed the procedures described above, except that the antibodies were cross-linked to protein A/G magnetic beads to reduce the background in Western blot. For the cross-linking, antibodies(Rabbit IgG, a-PHF21A) were reacted with Protein A/G beads (1:1 mixture) overnight at 4 °C, then cross-linked by 10 *μ*M DMP (Thermo 21667) in 0.2M sodium borate pH 9.0 for 30 min at RT. The reaction was quenched with 0.2 M Tris/Cl pH 8.0 at room temperature for 1.5 hrs, and the antibody-conjugated beads were washed with IP buffer and used for IP reaction. The following antibodies were used for Western blots: PHF21A (This study), LSD1(Abcam ab17721), CoREST (Abcam ab183711), HDAC2(Santa Cruz sc-7899).

### Immunofluorescence

Cells were fixed with 4% paraformaldehyde for 10 minutes at room temperature, permeabilized with 1% Triton X-100 for 5 minutes, and washed two times with PBS + 0.02% Triton X-100, and blocked with 10% FBS for 30 minutes. Cells were incubated with primary antibodies overnight at 4°C, washed three times with PBS + 0.02% Triton X-100, and then incubated with secondary antibodies for 1 hour at room temperature. Primary antibodies used were anti-strep tag (GenScript, A01732, 1/1000), anti-PHF21A (in house, generated as described previously^35^, 1/1000), anti-MAP2 (Millipore, AB5543, 1/1000). Cells were washed three times with PBS + 0.02% Triton X-100 and then mounted with ProLong Diamond Antifade Mountant (Invitrogen, P36961). Images were acquired with an Olympus BX61 microscope.

### Protein expression and purification

E.coli BL21-DE3 was transformed with pGEX-4T-1 PHF21A PHD/AT-hook fragments. 10 ml liquid cultures were grown overnight at 37°C in LB media and then transferred into 1 liter LB media the following day. Growth was monitored until OD600 = 0.6, then 0.25 mM IPTG was added to induce expression, and cells were incubated at 15°C for 16 hours. Then, E.coli cells were harvested, and the cell pellets were frozen at -80°C. Cells were lysed in Wash Buffer (20 mM HEPES-KOH at pH 7.2, 150 mM KCl, 0.05% NP-40, 10% glycerol) with protease inhibitors with no EDTA (Sigma S8830) and then sonicated for five minutes at 50% amplitude. Then, the soluble fraction was recovered after centrifugation at max speed for 15 minutes. The soluble fraction was incubated with a glutathione sepharose (Thermo 16101) for two hours. The sepharose beads were washed five times with Wash Buffer, and the bound proteins were eluted with Wash Buffer containing 10 mM reduced glutathione at pH 7.5. The eluates were then dialyzed against fresh wash buffer, and the GST tag was cleaved by thrombin. Free GST and thrombin were removed with a second glutathione column and a Ni-NTA column (Qiagen 30210) to remove thrombin. Purified protein concentration was assessed with a Bradford Assay, and stocks were aliquoted and stored at -80°C with 20% glycerol added.

To reconstitute PHF21A-LSD1-CoREST complexes, we used the Bac-to-Bac Baculovirus Expression System (Thermo 10359016) to grow baculoviruses and express proteins in High Five insect cells (Thermofisher). Sf21 cells were used for baculovirus propagation and Hi5 cells were used for protein expression. For the bipartite LC complex, the LSD1 N-terminus was MBP-tagged. For the tripartite PLC complex, the PHF21A N-terminus was fused to MBP. One subunit, either PHF21A or LSD1, was MBP-tagged at their N-termini. Following harvesting, High Five cells were lysed by homogenization in High Salt Wash Buffer (30 mM Tris/Cl at pH 7.5, 500 mM NaCl, 0.05% beta-mercaptoethanol) containing a protease inhibitor cocktail with no EDTA. Lysates were passed through a 26G needle twice and cleared with centrifugation at max speed for two hours. The soluble fraction was incubated with an amylose column (NEB E8021S) for two hours, and the amylose column was washed five times with High Salt Wash Buffer. Then the proteins were eluted with 10 mM maltose. The eluates were dialyzed against High Salt Wash Buffer and treated with TEV protease to remove the MBP tag. Then, purified complexes were subject to FPLC to separate the fraction with stoichiometric bipartite or tripartite subunit compositions.

Recombinant nucleosomes with bacterially expressed histones and 177-bp Widom601 DNA with were expressed and purified as previously described^64^. We also used recombinant designer nucleosomes carrying dimethylation at H3K4, H3K9, H3K27, H3K36, and H4K20 (EpiCypher). Cellular nucleosomes were isolated from HEK293T, stably expressing strep-tagged H2A at the N-terminus, which we generated with lentivirus. Strep-H2A-293T cells were incubated in hypotonic lysis buffer (20 mM HEPES-KOH at pH 7.2, 20 mM KCl, 10% glycerol, and 0.05% NP-40) with protease inhibitors for 10 minutes at 4°C. Then, chromatin was digested with Micrococcal Nuclease (NEB) in 20 mM HEPES-KOH at pH 7.2, 340 mM KCl, 10% glycerol, 0.05% NP-40, and 1mM CaCl2 for 2 hours. The reaction was quenched with 4 mM EGTA and centrifuged at max speed for 10 minutes to recover the solubilized chromatin. The soluble fraction was incubated with a Streptactin sepharose (Qiagen 1057979) for 3 hours, and the sepharose beads were washed five times with wash buffer (20 mM HEPES-KOH at pH 7.2, 150 mM KCl, 10% glycerol, 0.05% NP-40). Bound nucleosomes were eluted with wash buffer containing 50 mM desthiobiotin (IBA 2-1000-002) at pH 8.0. We confirmed ∼90% was mono-nucleosome by electrophoresis of DNA from the isolated nucleosome (data not shown).

### Electrophoretic mobility shift assays (EMSA)

100 nM Widom601 DNA or 1 *μ*g of nucleosomes were incubated with an increasing concentration of PHF21A protein fragments in EMSA Buffer (10 mM Tris/Cl at pH 7.5, 50 mM KCl, 0.5 mM MgCl2, 0.1 mM EDTA, 5% glycerol, and 0.1 mg/ml BSA) for 30 minutes at 25°C. Samples were loaded onto 6% polyacrylamide gels with 0.5X TBE, electrophoresed at 4 °C, and stained by ethidium bromide or SYBR Gold (Thermofisher).

### Histone-peptide binding assays

One *μ*g of histone H3 peptides with or without H3K4me2 (a.a. 1-21, Active Motif) were incubated with 1 *μ*g of purified GST-PHF21A protein fragments in binding buffer (50 mM Tris/Cl at pH 7.5, 150 mM NaCl, 0.05% NP-40) and rotated overnight. The following day, Streptavidin agarose beads (Thermo 20349) were incubated with the binding reaction for 1 hour at 4 °C to trap the peptide-PHF21A complex. Beads were washed five times with binding buffer and then denatured in Laemmli buffer, and the eluted proteins were subjected to Western Blot analysis using a GST antibody (1:1000, Millipore 05-782).

### Histone demethylation assays with reconstituted complexes

One *μ*g of recombinant (Epicypher) or cellular nucleosomes (see protein purification above) were incubated with 1, 3, or 9 *μ*g of LC or PLC complexes in histone demethylation buffer (50 mM Tris/Cl at pH 8.0, 50 mM KCl, 0.5% BSA, 5% glycerol) and incubated for 4 hours at 37°C. Then, samples went through SDS-PAGE and quantitative fluorescent Western Blot analysis (LI-COR). LI-COR Image Studio software was used to quantify the bands. The following antibodies were used: H4 (Abcam ab31830), H3 (Abcam ab24834), H3K4me1 (Abcam ab8895), H3K4me2 (Abcam ab7766), H3K9me1 (Epicypher 13-0029), H3K9me2 (Abcam ab194680), H3K27me1 (Abcam ab194688), H3K27me2 (Active Motif 39245), H3K36me1 (Abcam ab9048), H3K36me2 (Abcam ab9049), H4K20me1 (Abcam ab9051), H4K20me2 (Abcam ab9052). LiCOR Image Studio software was used to quantify histone modification band level relative to a histone H3 or H4 loading control. The averages of at least two independent experiments were reported.

### mini-gene splicing assays

Splicing assay constructs were generated by amplifying regions of genomic DNA containing the PHF21A alternative exons (exons 13-15 in RefSeq PHF21A transcript NM_001101802, where exon 14 is either the canonical or neuronal alternative exon). 200-300 base pairs of flanking intronic DNA were included to ensure the inclusion of splice branch sites. Specifically, the following three genomic regions were amplified and cloned by Gibson assembly: hg19 chr11: 45,970,763-45,970,212; 45,967,840-45,967,160; and 45,960,062-45,959,428. Point mutagenesis was used to introduce the patient mutation (1285 G>A) in exon 13. Following assembly, sequences were confirmed by Sanger sequencing and cloned into a pENTR plasmid. Gateway recombination was then used to generate plasmids with either a PGK or the SYN promoter and a Strep-tag.

PGK-promoter constructs were transfected into HEK293T cells using Lipofectamine 2000 according to standard protocols. One day post-transfection, cells were harvested in Trizol. hSYN1-promoter constructs were packaged into lentivirus and introduced into DIV3 mouse cortical neurons with MOI of 1. Neurons were harvested at DIV6 in Trizol. Total RNA was extracted with the Trizol-Chloroform method, followed by DNase treatment (Promega #M6101, US) and RNeasy Plus Mini kit (Qiagen Valencia, CA) for RNA cleanup following the manufacturer’s instructions. cDNA was synthesized using ProtoScript II Reverse Transcriptase (New England BioLab, US) according to the manufacturer’s protocol. All RT-qPCR reactions were run using Applied Biosystems™ Power SYBR™ Green PCR Master Mix (Thermo Fisher Scientific, US) and were analyzed (Applied Biosystems 7500 Instrument). The following primers were used: *PHF21A*-n (F1 and R2 in Figure 6B): F1 GTGGAGCCACCCCCAGT, R2 TTTGGCCAGTGTTCCTCATT. *PHF21A*-c (F1 and R3): F1 GTGGAGCCACCCCCAGT, R3 GCAGGGAAAGTGAATGTGGT. For the WPRE normalization (F4 and R5): F4 CGCTATGTGGATACGCTGCT, R5 GTTGCGTCAGCAAACACAGT. Splice products were gel extracted, Sanger sequenced, and confirmed as the spliced PHF21A mRNAs.

### Animals

All animal use followed NIH guidelines and was in compliance with the University of Michigan Committee on Use and Care of Animals.

## Notes

### Competing Interest Statement

The authors have declared no competing interest.

## References

1. Deplancke, B., Alpern, D., and Gardeux, V. (2016). The Genetics of Transcription Factor DNA Binding Variation. Cell 166, 538–554. 10.1016/j.cell.2016.07.012.

2. Zaret, K.S. (2018). Pioneering the chromatin landscape. Nature genetics 50, 167–169. 10.1038/s41588-017-0038-z.

3. Lorch, Y., LaPointe, J.W., and Kornberg, R.D. (1987). Nucleosomes inhibit the initiation of transcription but allow chain elongation with the displacement of histones. Cell 49, 203–210. 10.1016/0092-8674(87)90561-7.

4. Smith, E., and Shilatifard, A. (2010). The Chromatin Signaling Pathway: Diverse Mechanisms of Recruitment of Histone-Modifying Enzymes and Varied Biological Outcomes. Molecular Cell 40, 689–701. 10.1016/j.molcel.2010.11.031.

5. Deng, X., Qiu, Q., He, K., and Cao, X. (2018). The seekers: How epigenetic modifying enzymes find their hidden genomic targets in Arabidopsis. Current Opinion in Plant Biology 45, 75–81. 10.1016/j.pbi.2018.05.006.

6. Lessard, J., Wu, J.I., Ranish, J.A., Wan, M., Winslow, M.M., Staahl, B.T., Wu, H., Aebersold, R., Graef, I.A., and Crabtree, G.R. (2007). An essential switch in subunit composition of a chromatin remodeling complex during neural development. Neuron 55, 201–215. 10.1016/j.neuron.2007.06.019.

7. Olave, I., Wang, W., Xue, Y., Kuo, A., and Crabtree, G.R. (2002). Identification of a polymorphic, neuron-specific chromatin remodeling complex. Genes & development 16, 2509–2517. 10.1101/gad.992102.

8. Hota, S.K., Johnson, J.R., Verschueren, E., Thomas, R., Blotnick, A.M., Zhu, Y., Sun, X., Pennacchio, L.A., Krogan, N.J., and Bruneau, B.G. (2019). Dynamic BAF chromatin remodeling complex subunit inclusion promotes temporally distinct gene expression programs in cardiogenesis. Development 146. 10.1242/dev.174086.

9. Lickert, H., Takeuchi, J.K., Von Both, I., Walls, J.R., McAuliffe, F., Adamson, S.L., Henkelman, R.M., Wrana, J.L., Rossant, J., and Bruneau, B.G. (2004). Baf60c is essential for function of BAF chromatin remodelling complexes in heart development. Nature 432, 107–112. 10.1038/nature03071.

10. Goodrich, J.A., and Tjian, R. (2010). Unexpected roles for core promoter recognition factors in cell-type-specific transcription and gene regulation. Nat Rev Genet 11, 549–558. 10.1038/nrg2847.

11. Herrera, F.J., Yamaguchi, T., Roelink, H., and Tjian, R. (2014). Core promoter factor TAF9B regulates neuronal gene expression. Elife 3, e02559. 10.7554/eLife.02559.

12. Irimia, M., Weatheritt, R.J., Ellis, J.D., Parikshak, N.N., Gonatopoulos-Pournatzis, T., Babor, M., Quesnel-Vallieres, M., Tapial, J., Raj, B., O’Hanlon, D., et al. (2014). A highly conserved program of neuronal microexons is misregulated in autistic brains. Cell 159, 1511–1523. 10.1016/j.cell.2014.11.035.

13. Mele, M., Ferreira, P.G., Reverter, F., DeLuca, D.S., Monlong, J., Sammeth, M., Young, T.R., Goldmann, J.M., Pervouchine, D.D., Sullivan, T.J., et al. (2015). Human genomics. The human transcriptome across tissues and individuals. Science (New York, N.Y.) 348, 660– 665. 10.1126/science.aaa0355.

14. Yeo, G., Holste, D., Kreiman, G., and Burge, C.B. (2004). Variation in alternative splicing across human tissues. Genome biology5, R74. 10.1186/gb-2004-5-10-r74.

15. Makino, S., Kaji, R., Ando, S., Tomizawa, M., Yasuno, K., Goto, S., Matsumoto, S., Tabuena, M.D., Maranon, E., Dantes, M., et al. (2007). Reduced neuron-specific expression of the TAF1 gene is associated with X-linked dystonia-parkinsonism. American journal of human genetics 80, 393–406. 10.1086/512129.

16. Fiszbein, A., Giono, L.E., Quaglino, A., Berardino, B.G., Sigaut, L., Bilderling, C. von, Schor, I.E., Steinberg, J.H., Rossi, M., Pietrasanta, L.I., et al. (2016). Alternative Splicing of G9a Regulates Neuronal Differentiation. Cell Rep 14, 2797–2808. 10.1016/j.celrep.2016.02.063.

17. Zibetti, C., Adamo, A., Binda, C., Forneris, F., Toffolo, E., Verpelli, C., Ginelli, E., Mattevi, A., Sala, C., and Battaglioli, E. (2010). Alternative splicing of the histone demethylase LSD1/KDM1 contributes to the modulation of neurite morphogenesis in the mammalian nervous system. The Journal of neuroscience : the official journal of the Society for Neuroscience 30, 2521–2532. 10.1523/JNEUROSCI.5500-09.2010.

18. Shi, Y., Lan, F., Matson, C., Mulligan, P., Whetstine, J.R., Cole, P.A., Casero, R.A., and Shi, Y. (2004). Histone demethylation mediated by the nuclear amine oxidase homolog LSD1. Cell 119, 941–953. 10.1016/j.cell.2004.12.012.

19. Forneris, F., Binda, C., Vanoni, M.A., Mattevi, A., and Battaglioli, E. (2005). Histone demethylation catalysed by LSD1 is a flavin-dependent oxidative process. FEBS Lett 579, 2203–2207. 10.1016/j.febslet.2005.03.015.

20. Laurent, B., Ruitu, L., Murn, J., Hempel, K., Ferrao, R., Xiang, Y., Liu, S., Garcia, B.A., Wu, H., Wu, F., et al. (2015). A specific LSD1/KDM1A isoform regulates neuronal differentiation through H3K9 demethylation. Mol Cell 57, 957–970. 10.1016/j.molcel.2015.01.010.

21. Rusconi, F., Paganini, L., Braida, D., Ponzoni, L., Toffolo, E., Maroli, A., Landsberger, N., Bedogni, F., Turco, E., Pattini, L., et al. (2015). LSD1 Neurospecific Alternative Splicing Controls Neuronal Excitability in Mouse Models of Epilepsy. Cerebral cortex (New York, N.Y. : 1991) 25, 2729–2740. 10.1093/cercor/bhu070.

22. Rusconi, F., Grillo, B., Ponzoni, L., Bassani, S., Toffolo, E., Paganini, L., Mallei, A., Braida, D., Passafaro, M., Popoli, M., et al. (2016). LSD1 modulates stress-evoked transcription of immediate early genes and emotional behavior. Proceedings of the National Academy of Sciences of the United States of America 113, 3651–3656. 10.1073/pnas.1511974113.

23. Toffolo, E., Rusconi, F., Paganini, L., Tortorici, M., Pilotto, S., Heise, C., Verpelli, C., Tedeschi, G., Maffioli, E., Sala, C., et al. (2014). Phosphorylation of neuronal Lysine-Specific Demethylase 1LSD1/KDM1A impairs transcriptional repression by regulating interaction with CoREST and histone deacetylases HDAC1/2. Journal of neurochemistry 128, 603–616. 10.1111/jnc.12457.

24. Wang, J., Telese, F., Tan, Y., Li, W., Jin, C., He, X., Basnet, H., Ma, Q., Merkurjev, D., Zhu, X., et al. (2015). LSD1n is an H4K20 demethylase regulating memory formation via transcriptional elongation control. Nature neuroscience 18, 1256–1264. 10.1038/nn.4069.

25. Medvedeva, Y.A., Lennartsson, A., Ehsani, R., Kulakovskiy, I.V., Vorontsov, I.E., Panahandeh, P., Khimulya, G., Kasukawa, T., Consortium, F., and Drablos, F. (2015). EpiFactors: A comprehensive database of human epigenetic factors and complexes. Database : the journal of biological databases and curation 2015, bav067. 10.1093/database/bav067.

26. Porter, R.S., Jaamour, F., and Iwase, S. (2018). Neuron-specific alternative splicing of transcriptional machineries: Implications for neurodevelopmental disorders. Molecular and cellular neurosciences 87, 35–45. 10.1016/j.mcn.2017.10.006.

27. Zhang, Y., Chen, K., Sloan, S.A., Bennett, M.L., Scholze, A.R., O’Keeffe, S., Phatnani, H.P., Guarnieri, P., Caneda, C., Ruderisch, N., et al. (2014). An RNA-sequencing transcriptome and splicing database of glia, neurons, and vascular cells of the cerebral cortex. The Journal of neuroscience : the official journal of the Society for Neuroscience 34, 11929–11947. 10.1523/JNEUROSCI.1860-14.2014.

28. Consortium, G. (2015). Human genomics. The Genotype-Tissue Expression (GTEx) pilot analysis: Multitissue gene regulation in humans. Science 348, 648–660. 10.1126/science.1262110.

29. Hakimi, M.A., Bochar, D.A., Chenoweth, J., Lane, W.S., Mandel, G., and Shiekhattar, R. (2002). A core-BRAF35 complex containing histone deacetylase mediates repression of neuronal-specific genes. Proceedings of the National Academy of Sciences of the United States of America 99, 7420–7425. 10.1073/pnas.112008599.

30. Shi, Y.J., Matson, C., Lan, F., Iwase, S., Baba, T., and Shi, Y. (2005). Regulation of LSD1 histone demethylase activity by its associated factors. Molecular cell 19, 857–864. 10.1016/j.molcel.2005.08.027.

31. Lan, F., Collins, R.E., De Cegli, R., Alpatov, R., Horton, J.R., Shi, X., Gozani, O., Cheng, X., and Shi, Y. (2007). Recognition of unmethylated histone H3 lysine 4 links BHC80 to LSD1-mediated gene repression. Nature 448, 718–722. 10.1038/nature06034.

32. Andres, M.E., Burger, C., Peral-Rubio, M.J., Battaglioli, E., Anderson, M.E., Grimes, J., Dallman, J., Ballas, N., and Mandel, G. (1999). CoREST: A functional corepressor required for regulation of neural-specific gene expression. Proceedings of the National Academy of Sciences of the United States of America 96, 9873–9878.

33. Lee, M.G., Wynder, C., Cooch, N., and Shiekhattar, R. (2005). An essential role for CoREST in nucleosomal histone 3 lysine 4 demethylation. Nature 437, 432–435. 10.1038/nature04021.

34. Yang, M., Gocke, C.B., Luo, X., Borek, D., Tomchick, D.R., Machius, M., Otwinowski, Z., and Yu, H. (2006). Structural basis for CoREST-dependent demethylation of nucleosomes by the human LSD1 histone demethylase. Molecular cell 23, 377–387. 10.1016/j.molcel.2006.07.012.

35. Iwase, S., Januma, A., Miyamoto, K., Shono, N., Honda, A., Yanagisawa, J., and Baba, T. (2004). Characterization of BHC80 in BRAF-HDAC complex, involved in neuron-specific gene repression. Biochemical and biophysical research communications 322, 601–608. 10.1016/j.bbrc.2004.07.163.

36. Furlanis, E., Traunmuller, L., Fucile, G., and Scheiffele, P. (2019). Landscape of ribosome-engaged transcript isoforms reveals extensive neuronal-cell-class-specific alternative splicing programs. Nat Neurosci 22, 1709–1717. 10.1038/s41593-019-0465-5.

37. Li, W., Sun, W., Zhang, Y., Wei, W., Ambasudhan, R., Xia, P., Talantova, M., Lin, T., Kim, J., Wang, X., et al. (2011). Rapid induction and long-term self-renewal of primitive neural precursors from human embryonic stem cells by small molecule inhibitors. Proceedings of the National Academy of Sciences 108, 8299–8304. 10.1073/pnas.1014041108.

38. Brennand, K.J., Simone, A., Jou, J., Gelboin-Burkhart, C., Tran, N., Sangar, S., Li, Y., Mu, Y., Chen, G., Yu, D., et al. (2011). Modelling schizophrenia using human induced pluripotent stem cells. Nature 473, 221–225. 10.1038/nature09915.

39. Utami, K.H., Skotte, N.H., Colaço, A.R., Yusof, N.A.B.M., Sim, B., Yeo, X.Y., Bae, H.-G., Garcia-Miralles, M., Radulescu, C.I., Chen, Q., et al. (2020). Integrative Analysis Identifies Key Molecular Signatures Underlying Neurodevelopmental Deficits in Fragile X Syndrome. Biological Psychiatry 88, 500–511. 10.1016/j.biopsych.2020.05.005.

40. Busskamp, V., Lewis, N.E., Guye, P., Ng, A.H., Shipman, S.L., Byrne, S.M., Sanjana, N.E., Murn, J., Li, Y., Li, S., et al. (2014). Rapid neurogenesis through transcriptional activation in human stem cells. Molecular Systems Biology 10, 760. 10.15252/msb.20145508.

41. Li, Y., Xie, N., Chen, R., Lee, A.R., Lovnicki, J., Morrison, E.A., Fazli, L., Zhang, Q., Musselman, C.A., Wang, Y., et al. (2019). RNA Splicing of the BHC80 Gene Contributes to Neuroendocrine Prostate Cancer Progression. European Urology 76, 157–166. 10.1016/j.eururo.2019.03.011.

42. Reeves, R., and Nissen, M.S. (1990). The A.T-DNA-binding domain of mammalian high mobility group I chromosomal proteins. A novel peptide motif for recognizing DNA structure. The Journal of biological chemistry 265, 8573–8582.

43. Kim, S.A., Zhu, J., Yennawar, N., Eek, P., and Tan, S. (2020). Crystal Structure of the LSD1/CoREST Histone Demethylase Bound to Its Nucleosome Substrate. Molecular cell 78, 903–914 e4. 10.1016/j.molcel.2020.04.019.

44. Kim, H.G., Kim, H.T., Leach, N.T., Lan, F., Ullmann, R., Silahtaroglu, A., Kurth, I., Nowka, A., Seong, I.S., Shen, Y., et al. (2012). Translocations disrupting PHF21A in the Potocki-Shaffer-syndrome region are associated with intellectual disability and craniofacial anomalies. Am J Hum Genet 91, 56–72. 10.1016/j.ajhg.2012.05.005.

45. Kim, H.G., Rosenfeld, J.A., Scott, D.A., Benedicte, G., Labonne, J.D., Brown, J., McGuire, M., Mahida, S., Naidu, S., Gutierrez, J., et al. (2019). Disruption of PHF21A causes syndromic intellectual disability with craniofacial anomalies, epilepsy, hypotonia, and neurobehavioral problems including autism. Molecular autism 10, 35. 10.1186/s13229-019-0286-0.

46. Labonne, J.D., Vogt, J., Reali, L., Kong, I.K., Layman, L.C., and Kim, H.G. (2015). A microdeletion encompassing PHF21A in an individual with global developmental delay and craniofacial anomalies. American journal of medical genetics. Part A 167A, 3011–3018. 10.1002/ajmg.a.37344.

47. Potocki, L., and Shaffer, L.G. (1996). Interstitial deletion of 11(p11.2p12): A newly described contiguous gene deletion syndrome involving the gene for hereditary multiple exostoses (EXT2). American journal of medical genetics 62, 319–325. 10.1002/(sici)1096-8628(19960329)62:3<319::aid-ajmg22>3.0.co;2-m.

48. Satterstrom, F.K., Kosmicki, J.A., Wang, J., Breen, M.S., De Rubeis, S., An, J.Y., Peng, M., Collins, R., Grove, J., Klei, L., et al. (2020). Large-Scale Exome Sequencing Study Implicates Both Developmental and Functional Changes in the Neurobiology of Autism. Cell 180, 568– 584 e23. 10.1016/j.cell.2019.12.036.

49. Stroud, H., Su, S.C., Hrvatin, S., Greben, A.W., Renthal, W., Boxer, L.D., Nagy, M.A., Hochbaum, D.R., Kinde, B., Gabel, H.W., et al. (2017). Early-Life Gene Expression in Neurons Modulates Lasting Epigenetic States. Cell 171, 1151–1164 e16. 10.1016/j.cell.2017.09.047.

50. Clark, S.C., Chereji, R.V., Lee, P.R., Fields, R.D., and Clark, D.J. (2020). Differential nucleosome spacing in neurons and glia. Neurosci Lett 714, 134559. 10.1016/j.neulet.2019.134559.

51. Kadoch, C., and Crabtree, G.R. (2015). Mammalian SWI/SNF chromatin remodeling complexes and cancer: Mechanistic insights gained from human genomics. Sci Adv 1, e1500447. 10.1126/sciadv.1500447.

52. De Rubeis, S., He, X., Goldberg, A.P., Poultney, C.S., Samocha, K., Cicek, A.E., Kou, Y., Liu, L., Fromer, M., Walker, S., et al. (2014). Synaptic, transcriptional and chromatin genes disrupted in autism. Nature 515, 209–215. 10.1038/nature13772.

53. Iossifov, I., O’Roak, B.J., Sanders, S.J., Ronemus, M., Krumm, N., Levy, D., Stessman, H.A., Witherspoon, K.T., Vives, L., Patterson, K.E., et al. (2014). The contribution of de novo coding mutations to autism spectrum disorder. Nature 515, 216–221. 10.1038/nature13908.

54. Najmabadi, H., Hu, H., Garshasbi, M., Zemojtel, T., Abedini, S.S., Chen, W., Hosseini, M., Behjati, F., Haas, S., Jamali, P., et al. (2011). Deep sequencing reveals 50 novel genes for recessive cognitive disorders. Nature 478, 57–63. 10.1038/nature10423.

55. Network, and Pathway Analysis Subgroup of Psychiatric Genomics, C. (2015). Psychiatric genome-wide association study analyses implicate neuronal, immune and histone pathways. Nature neuroscience 18, 199–209. 10.1038/nn.3922.

56. Huth, J.R., Bewley, C.A., Nissen, M.S., Evans, J.N., Reeves, R., Gronenborn, A.M., and Clore, G.M. (1997). The solution structure of an HMG-I(Y)-DNA complex defines a new architectural minor groove binding motif. Nat. Struct. Biol. 4, 657–665.

57. Fonfría-Subirós, E., Acosta-Reyes, F., Saperas, N., Pous, J., Subirana, J.A., and Campos, J.L. (2012). Crystal structure of a complex of DNA with one AT-hook of HMGA1. PLoS ONE 7, e37120. 10.1371/journal.pone.0037120.

58. Geierstanger, B.H., Volkman, B.F., Kremer, W., and Wemmer, D.E. (1994). Short peptide fragments derived from HMG-i/y proteins bind specifically to the minor groove of DNA. Biochemistry 33, 5347–5355. 10.1021/bi00183a043.

59. Xue, Y., Ouyang, K., Huang, J., Zhou, Y., Ouyang, H., Li, H., Wang, G., Wu, Q., Wei, C., Bi, Y., et al. (2013). Direct Conversion of Fibroblasts to Neurons by Reprogramming PTB-Regulated MicroRNA Circuits. Cell 152, 82–96. 10.1016/j.cell.2012.11.045.

60. Raj, B., Irimia, M., Braunschweig, U., Sterne-Weiler, T., O’Hanlon, D., Lin, Z.-Y., Chen, G.I., Easton, L.E., Ule, J., Gingras, A.-C., et al. (2014). A Global Regulatory Mechanism for Activating an Exon Network Required for Neurogenesis. Molecular Cell 56, 90–103. 10.1016/j.molcel.2014.08.011.

61. Marchler-Bauer, A., Bo, Y., Han, L., He, J., Lanczycki, C.J., Lu, S., Chitsaz, F., Derbyshire, M.K., Geer, R.C., Gonzales, N.R., et al. (2017). CDD/SPARCLE: Functional classification of proteins via subfamily domain architectures. Nucleic acids research 45, D200–d203. 10.1093/nar/gkw1129.

62. Doyle, S.A. (2005). High-throughput cloning for proteomics research. Methods Mol Biol 310, 107–113. 10.1007/978-1-59259-948-6_7.

63. Garay, P.M., Chen, A., Tsukahara, T., Rodriguez Diaz, J.C., Kohen, R., Althaus, J.C., Wallner, M.A., Giger, R.J., Jones, K.S., Sutton, M.A., et al. (2020). RAI1 Regulates Activity-Dependent Nascent Transcription and Synaptic Scaling. Cell reports 32, 108002. 10.1016/j.celrep.2020.108002.

64. Lee, Y.T., Gibbons, G., Lee, S.Y., Nikolovska-Coleska, Z., and Dou, Y. (2015). One-pot refolding of core histones from bacterial inclusion bodies allows rapid reconstitution of histone octamer. Protein Expr Purif 110, 89–94. 10.1016/j.pep.2015.02.007.

